# Single-cell optogenetics reveals attenuation-by-suppression in visual cortical neurons

**DOI:** 10.1101/2023.09.13.557650

**Authors:** Paul K. LaFosse, Zhishang Zhou, Jonathan F. O’Rawe, Nina G. Friedman, Victoria M. Scott, Yanting Deng, Mark H. Histed

## Abstract

The relationship between neurons’ input and spiking output is central to brain computation. Studies *in vitro* and in anesthetized animals suggest nonlinearities emerge in cells’ input-output (activation) functions as network activity increases, yet how neurons transform inputs *in vivo* has been unclear. Here, we characterize cortical principal neurons’ activation functions in awake mice using two-photon optogenetics. We deliver fixed inputs at the soma while neurons’ activity varies with sensory stimuli. We find responses to fixed optogenetic input are nearly unchanged as neurons are excited, reflecting a linear response regime above neurons’ resting point. In contrast, responses are dramatically attenuated by suppression. This attenuation is a powerful means to filter inputs arriving to suppressed cells, privileging other inputs arriving to excited neurons. These results have two major implications. First, somatic neural activation functions *in vivo* accord with the activation functions used in recent machine learning systems. Second, neurons’ IO functions can filter sensory inputs — not only do sensory stimuli change neurons’ spiking outputs, but these changes also affect responses to input, attenuating responses to some inputs while leaving others unchanged.

**Significance statement:** How neurons transform their inputs into outputs is a fundamental building block of brain computation. Past studies have measured neurons’ input-output (IO) functions *in vitro* or in anesthetized states. Here, we measure neurons’ IO functions in the awake and intact brain, where ongoing network activity can influence neurons’ responses to input. Using state-of-the-art optogenetic methods to deliver precise inputs to neurons near the cell body, or soma, we discover neurons have a supralinear-to-linear IO function, contrary to previous findings of threshold-linear, strongly saturating, or power law IO functions. This supralinear-to-linear somatic IO function shape allows neurons to decrease their responses to, or filter, inputs while they are suppressed below their resting firing rates, a computation we term attenuation-by-suppression.

## Introduction

Neurons transform their synaptic inputs into output activity. The way in which neurons, and connected networks of neurons, achieve these transformations is a central and critical component of neural computation. For single neurons, frequency-current (f-I) curves describe how aggregate synaptic input generates spiking. F-I curves (in different contexts called static nonlinearities, neural transfer functions (McCormick et al., 1985; Powers and Binder, 2001; Fourcaud-Trocmé et al., 2003), or what we will use in this work, input-output or IO functions) thus govern how spikes are produced and passed down to postsynaptic partners. At the network level, the shape of the input-output function affects performance of the overall network. For example, IO function shape affects how well sensory networks can encode features of the world (Moskovitz et al., 2018). Machine learning (ML) work has confirmed that the shape of IO functions impacts network function, in part by exerting influence on learning speed and accuracy. For example, now-classic work has found that rectified-linear (ReLU) IO functions (or activation functions, the name often used in the ML context) can make learning more efficient and robust than learning with sigmoidal activation functions (Glorot et al., 2011; Nair and Hinton, 2010; Krizhevsky et al., 2012).

While work in all these fields often assumes for simplicity that neural IO functions are fixed or static, in real biological networks IO function shape is critically affected by all the inputs neurons receive — that is, IO function shape is affected by activity in the local recurrent network (Chance et al., 2002; Hansel and van Vreeswijk, 2002; Miller and Troyer, 2002; Priebe and Ferster, 2002; Agüera y Arcas and Fairhall, 2003; Rauch et al., 2003; Carandini, 2004; Ostojic and Brunel, 2011; Atallah et al., 2012; Haider et al., 2013). Thus, accurately determining neural IO functions requires measurements *in vivo* during normal network operation.

Past work has reported various shapes for neural IO functions. *In vitro* studies, where network activity levels can be low, have often found ReLU (also called threshold-linear) functions (Granit et al., 1963; McCormick et al., 1985; Jagadeesh et al., 1992; Carandini et al., 1996; Ahmed et al., 1998; Chance et al., 2002). Studies done in networks with higher levels of network activity than expected *in vitro* — in anesthetized animals, *ex vivo*, or *in vitro* with simulated noise — have found power law functions, with a supralinear regime throughout the input range (Anzai et al., 1999; Anderson et al., 2000; Priebe et al., 2004; Finn et al., 2007; Cardin et al., 2008; Petersen and Berg, 2016). On the other hand, theoretical studies of integrate-and-fire networks of active neurons have found an analytical description of the IO function in those networks, the Ricciardi input-output function, which has a supralinear regime followed by a linear, then a sublinear regime (Ricciardi and Smith, 1977; Sanzeni et al., 2020).

Those different descriptions conflict in important ways, and therefore the IO function shape seen *in vivo* during awake states of normal operation has been unclear (Yamashita et al., 2013; Tan et al., 2014; Zhao et al., 2016; Gajowa, 2018). This conflict arises in part because electrophysiological studies with control of neural input are difficult to perform in awake animals (Dubois, 2000; Bar-Yehuda and Korngreen, 2008; Wilson et al., 2011). Here, we use two-photon optogenetics to expand measurements of IO functions to hundreds of neurons in awake mice. We deliver the same optogenetic input to single cells at different activity levels, and thus determine neurons’ input-output responses at different points along their IO function. We deliver input to the soma of cells using a somatically targeted opsin. Thus, here we study the somatic IO function —how aggregated inputs from the dendrites, or dendritic outputs, affect spike rates.

We use this approach to explore the somatic IO functions of neurons in visual cortex. We stimulate selected neurons optogenetically, deliver visual stimuli that excite or suppress cells, and measure differences in response to the same optogenetic stimuli. We find neurons *in vivo* have an IO function that resembles not a ReLU or a power law function, but the Ricciardi function that is created by integrate-and-fire networks. The measured IO functions, as input varies from low to high, have a supralinearity followed by a linear region, and show some saturation at the upper end of the range of normal input. That upper range is defined by normal network activation levels —neurons’ firing in response to strong, high-contrast visual stimuli. A new and key observation of this work is that neurons rest near the top of the supralinear region. This means suppressed neurons produce smaller responses to the same input, suggesting cortical neurons attenuate responses to input that arrive to suppressed cells. This attenuation-by-suppression may be a computational means by which single neurons filter or mask out certain inputs.

## Results

### Uncovering input-output nonlinearities via cell-specific optogenetic stimulation

We use cell-specific two-photon optogenetic stimulation (Marshel et al., 2019; Dalgleish et al., 2020; LaFosse et al., 2023) to measure cells’ IO functions. By providing the same fixed input at different points along a cells’ IO function, the corresponding evoked changes in output can be used to infer the IO function shape (Fig. 1A). For example, if a cell’s IO function were threshold-linear (Fig. 1B, left), the response to a fixed optogenetic input would remain unchanged as the cell’s activity level increases. Similarly, when that cell is suppressed below the threshold, input would elicit no response. Our stimulation bypasses dendritic nonlinearities, by delivering input to the cell at the soma (using a soma-targeted opsin, stChrimsonR), allowing us to measure the transformation of the cell’s aggregate input into output spiking.

**Fig. 1:**
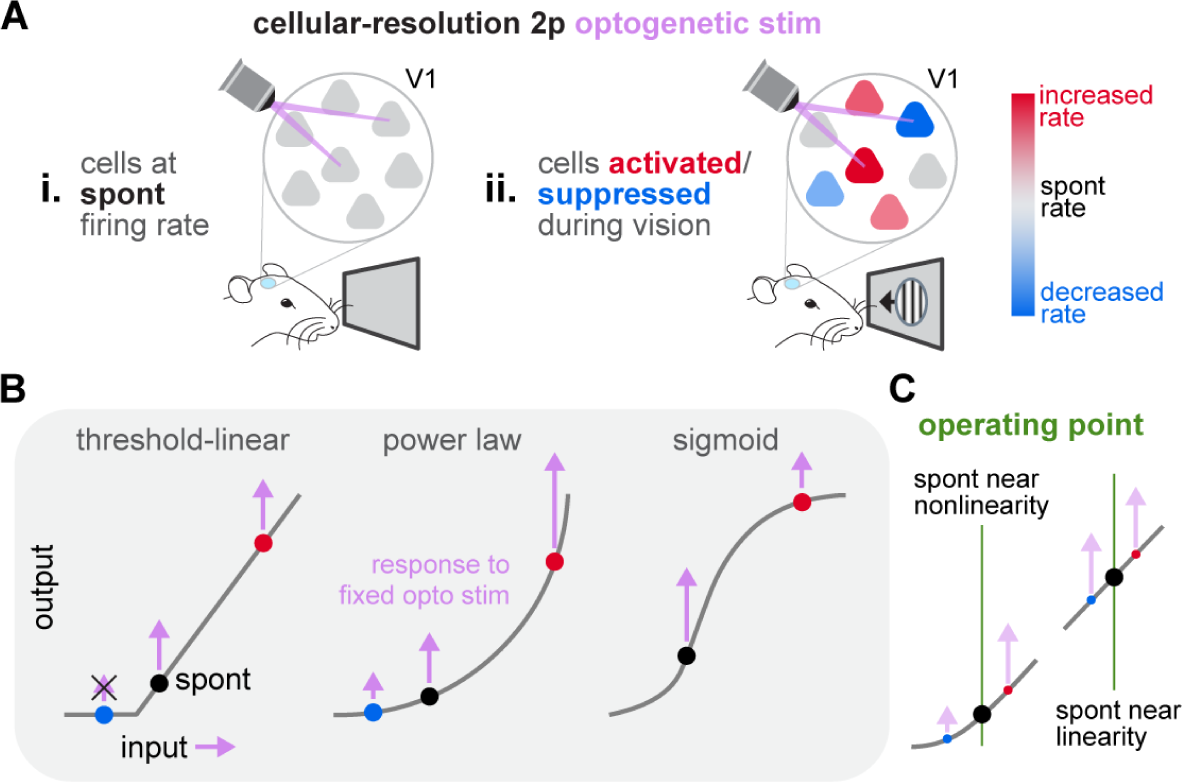
Mapping neurons’ input-output functions in awake mice using cellular-resolution optogenetic stimulation. **(A)** Schematic of experiments in V1 of awake mice during (**i**) spontaneous activity and (**ii**) visual stimulus presentation. **(B)** Schematics of some possible input-output (IO) functions. Optogenetic responses at different points along the IO function can uncover the shape and extent of nonlinearities. **(C)** Beyond the shape of the IO function, a key unknown feature is where neurons sit on their IO curves during spontaneous activity (schematized by vertical green lines).

The paper proceeds as follows. We deliver visual stimulation to change cells’ activity levels. We first deliver optogenetic stimuli paired with high-contrast visual stimuli that produce a wide range of responses, both suppression and activation, across populations of neurons in V1 (Fig. 2). Using these data we infer the shape of the average IO function across cells. We then show that this average shape is consistent with that seen in individual cells (Fig. 3). To do this we vary first visual stimuli to alternately suppress and activate the same cell, and then vary optogenetic stimulation intensity. We end by reconstructing this IO function, inferring the average operating point (Fig. 4; see Fig. 1C for a schematic), and discussing the major features of the IO function that we have uncovered.

**Fig. 2:**
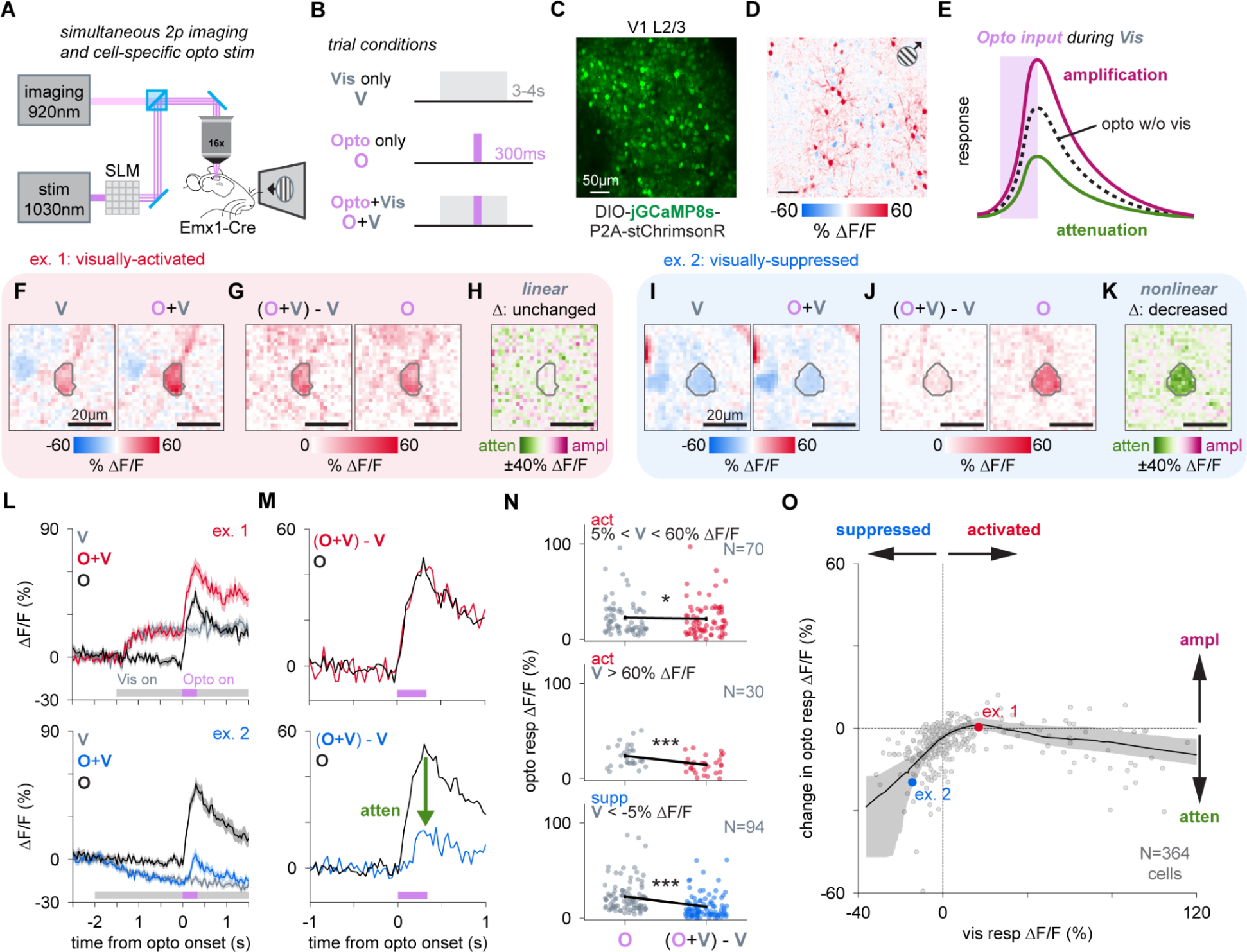
V1 neurons attenuate inputs when suppressed. **(A-B)** Experiment schematic. Two-photon calcium imaging during visual, cell-specific optogenetic, or paired optogenetic + visual stimulation in awake mice. **(C)** DIO-jGCaMP8s-P2A-stChrimsonR expressing in V1 layer 2/3. **(D)** Visually-evoked responses to a drifting grating: suppressed (blue) and activated (red) neurons. **(E)** Schematic showing increases/decreases of optogenetic responses during vision reveal nonlinear amplification/attenuation of inputs. **(F)** Example visually-activated cell. Visual response (left) and paired Opto+Vis response (right). **(G)** Optogenetic response during vision (O+V, left) and Opto only response (right). **(H)** No change in optogenetic response during and without vision (linear regime). **(I-K)** Same as **F-H** for a visually-suppressed cell with decreased optogenetic response during vision (nonlinear regime). **(L)** Trial-averaged (mean ± SEM, N = 50 reps) activity traces for the same visually-activated (top) and visually-suppressed (bottom) example cells in **F-K**. **(M)**, Comparison of optogenetic responses during and without vision. Green arrow depicts attenuation in visually-suppressed cell. **(N)** Visually-activated cells (visual response between 5% and 60% ΔF/F_0_; N = 70) show minimal change in optogenetic response (top). Strongly-activated cells (visual response > 60% ΔF/F_0_; N = 30) show decreased optogenetic response, suggesting saturation (middle). Visually-suppressed cells (visual response < -5% ΔF/F_0_; N = 94) show attenuated optogenetic response (bottom). *p < 0.05, ***p < 0.001; Wilcoxon signed-rank test with Bonferroni correction for multiple comparisons. Black lines: mean ± SEM. **(O)** Change in optogenetic response in neurons (N = 364 total cells, 15 ± 1.3 cells responsive per pattern; N = 25 patterns across N = 7 animals) across the working range of visual responses. Black line: LOESS fit to data with 95% confidence intervals via bootstrap.

**Fig. 3:**
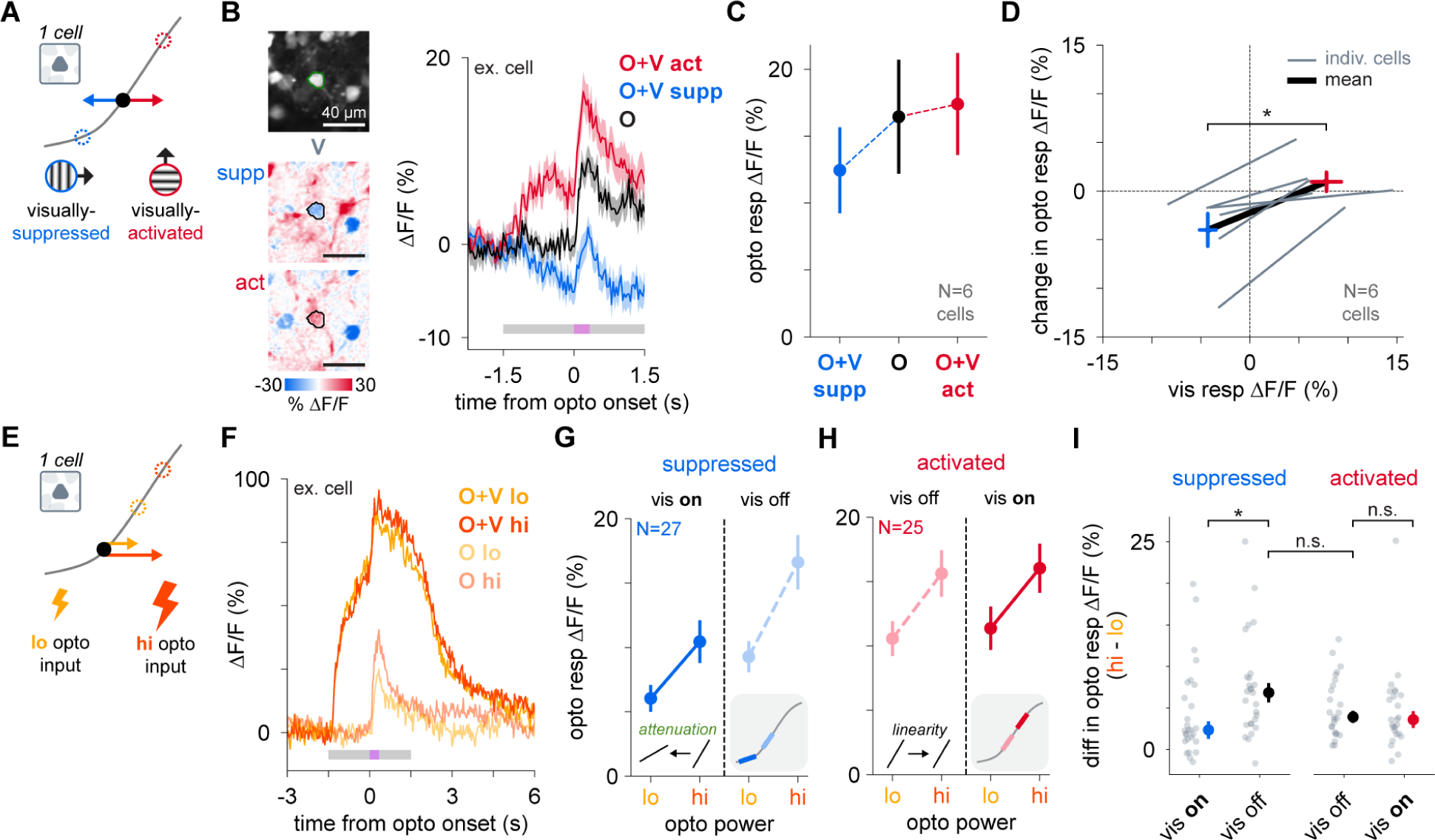
Individual V1 neurons exhibit nonlinear responses when suppressed and linear responses when activated. **(A)** Experiment schematic showing single cells suppressed and activated by orthogonal gratings, moving them down and up along their IO function. **(B)** Example neuron suppressed and activated by two different visual stimuli (left). Trial averaged traces (mean ± SEM; N = 50 trials) in Opto only and Opto+Vis conditions for both visual stimuli. **(C)** Cell-averaged (mean ± SEM; N = 6) optogenetic responses during and without visual stimuli. **(D)** Change in optogenetic response plotted against visual response for all cells. Gray lines: individual cells at their suppressed and activated states. Black line: mean across cells. Crosses are SEM of change in optogenetic response and visual response across cells. Blue: suppressed. Red: activated. *p < 0.05; Wilcoxon signed-rank test. **(E)** Experiment schematic showing single cells stimulated with two optogenetic powers (lo: 6.5 mW/target, hi: 8 mW/target). **(F)** Example neuron stimulated with both optogenetic powers during and without visual stimulation. **(G)** Cell-averaged (mean ± SEM; N = 27) optogenetic responses in visually-suppressed cells for vis on and vis off conditions. Inset: as cells are visually-suppressed, they move to a lower slope region of their IO function, measured between the two points of optogenetic powers. **(H)** Same as **G**, for visually-activated cells (N = 25). Inset: as cells are visually-activated, they remain in a linear regime of their IO function. **(I)** Difference in optogenetic response between low and high optogenetic powers. Grey points: individual cells. Median ± SEM across cells plotted for vis on (blue: suppressed, red: activated) and vis off (black) conditions. Suppressed cells show decreases in response difference between optogenetic powers. *p < 0.05; Wilcoxon signed-rank test (vis on vs. vis off) or Mann-Whitney U rank test (vis off vs. vis off) with Bonferroni correction for multiple comparisons.

**Fig. 4:**
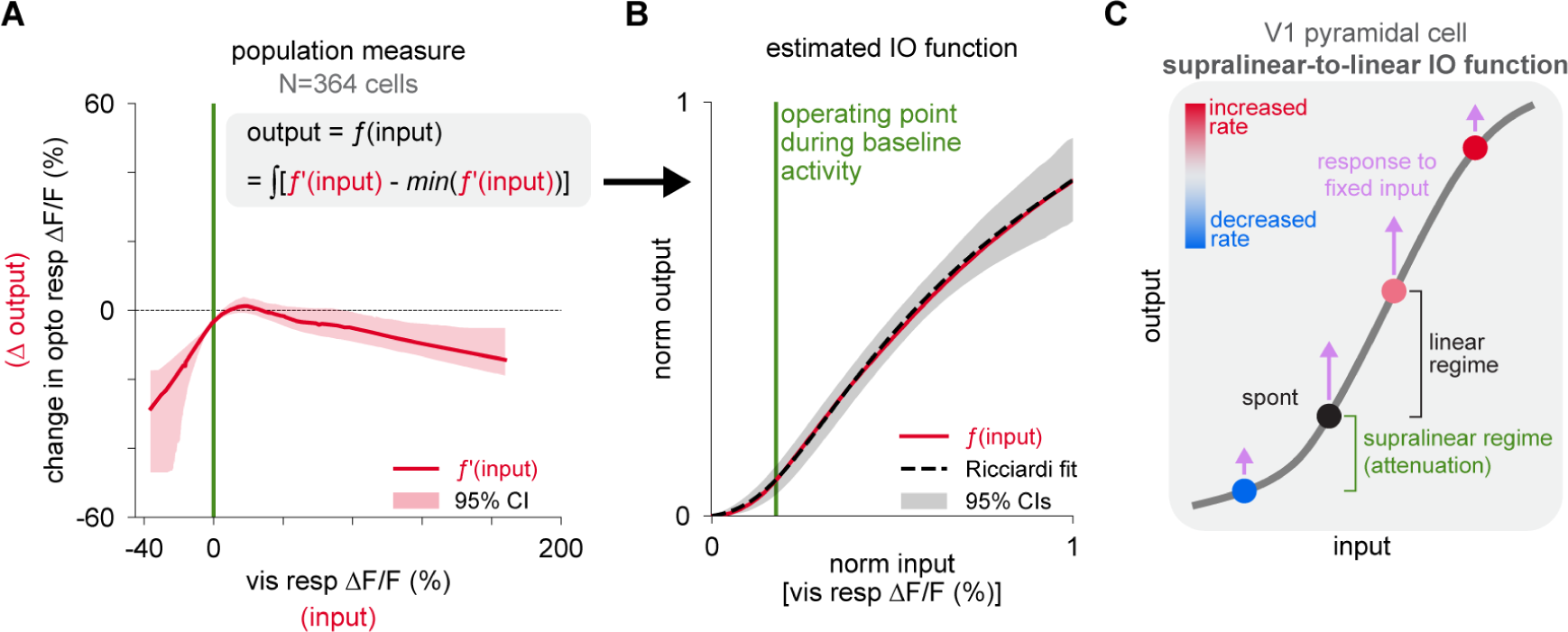
V1 pyramidal cells operate at the transition point of a supralinear-to-linear IO function. **(A)** Change in optogenetic response as a function of visual response. Red line: LOESS fit to cell data (N = 364 cells) with 95% confidence interval. **(B)** Estimated IO function and 95% confidence intervals calculated by shifting and integrating the fit trend line in **A** (see Methods). Green line: the operating point of V1 cells, set as the point in which cells’ visual response is 0% ΔF/F_0_ (green line in **A**). Black line: Ricciardi transfer function fit to the estimated IO function (± 95% CIs via bootstrap; Methods). **(C)** Schematic of the underlying IO function of the average V1 pyramidal cell showing a supralinear regime for low inputs and a linear regime before response saturation for high inputs. Cells operate between the supralinear and linear regimes, allowing cells suppressed below their spontaneous activity levels to attenuate inputs.

### Cell-specific optogenetic stimulation during and without visual stimulation reveals attenuation of inputs with suppression

We expressed the soma-targeted opsin stChrimsonR and the calcium indicator jGCaMP8s via a single Cre-dependent virus (in the Emx1-Cre mouse line), and stimulated V1 excitatory pyramidal cells (Fig. 2A) at cellular resolution while imaging them. To compare responses to optogenetic stimulation at different points along cells’ IO functions, we stimulated during both spontaneous conditions (i.e., no concurrent sensory stimulation), as well as during presentation of a visual stimulus (Fig. 2B; trial types randomly interleaved; Methods). Many cells change their activity in response to drifting gratings, with many increasing their firing rates (visually-activated) and some decreasing (visually-suppressed; Fig. 2C-D; Fig. S1).

We applied optogenetic stimulation while cells’ firing rates were changed during visual stimulation, as well as during periods of spontaneous activity with no visual stimulus (visual stimulus: 3-4 seconds, optogenetic stimulus: 300 ms, with onset 1.5-2.0 s after visual stimulation onset; Methods). We compared the response produced by optogenetic stimulation during the visual stimulus (Opto+Vis, Fig. 2F; incremental response during visual stimulation defined as (Opto+Vis)-Vis, Fig. 2G) to the response during optogenetic stimulus alone (Opto, Fig. 2G).

If the IO function follows a power law (is concave up; Fig. 1B), the optogenetic response should increase as neurons’ activity increases (schematic: Fig. 2E). If cells instead operate in a linear regime, they should produce a similar response increment to optogenetic input even as neurons’ activity varies.

For many neurons whose firing rates were increased by visual stimulation, the response to stimulation was similar when optogenetic stimulation was delivered alone or with visual stimulation (example cell, Fig. 2H; population, Fig. 2N). Neurons that were moderately activated by the visual stimulus (N = 70 cells; ΔF/F_0_ > 5% and < 60%) showed nearly the same response in the two conditions (mean optogenetic response during spontaneous condition: 22.7% ΔF/F_0_, paired visual condition: 21.3%; p = 0.02, Wilcoxon signed-rank test with Bonferroni correction for multiple comparisons; Fig. 2N, top). This indicates that above neurons’ firing rate at rest, neurons’ average IO function is nearly linear.

In cells exhibiting large increases in firing rate, we found some signs of saturation. Neurons more strongly activated by the visual stimulus (N = 30 cells; ΔF/F_0_ > 60%) showed decreased responses to optogenetic input (p = 2*10^-4^, Wilcoxon signed-rank test with Bonferroni correction; Fig. 2N, middle).

In contrast to the linear effects we observed for increases in firing rate, many cells suppressed by visual stimulation showed substantial attenuation of their response to optogenetic input (Fig. 2I-K). The attenuation was often dramatic, as in the example cell (Fig. 2L-M, bottom) whose response was reduced to less than 50% of its response before suppression. Attenuation was widespread across the population of suppressed neurons (N = 94 suppressed cells, ΔF/F_0_ < -5%, attenuation average: 47.3%; p = 2*10^-16^, Wilcoxon signed-rank test with Bonferroni correction; Fig. 2N, bottom). These observations suggest that below neurons’ resting activity levels (below spontaneous activity) the IO function shape is not linear, and instead shows a substantial nonlinearity.

The timecourses of neurons’ responses to optogenetic input (Fig. 2L-M) were as expected for both suppressed and activated neurons, featuring a fast increase in fluorescence during optogenetic stimulation consistent with induced spiking, followed by a slower decay after optogenetic stimulus offset consistent with calcium decay dynamics. Across trials, visual responses remained relatively consistent, with minimal response adaptation. Optogenetic responses slowly decreased across trials, but our results were consistent when analysis was restricted to either the early or the late trials (Fig. S2).

To characterize neurons’ IO functions across a range of activity levels, we plotted responses across many optogenetic stimulation sessions in many cells, spanning the working range of visual responses — from strongly suppressed to strongly activated (N = 11 experiments in N = 7 mice; N = 364 total cells stimulated). We plotted the average change in optogenetic response as a function of the neurons’ responses to visual input just before receiving optogenetic stimulation (Fig. 2O). As cells were more suppressed, we found, on average, more attenuation (via LOESS regression; Methods). This attenuation was seen across individual animals and experiments (Fig. S3).

The effects did not depend on viral preparation (bicistronic virus or separate viruses with opsin and GCaMP; Fig. S4). The effects also did not co-vary with baseline fluorescence across cells or animals, suggesting that the results are not changed by differences in GCaMP expression (Fig. S4).

The results also did not change when the visual stimulus was varied. Neurons showed the same attenuation-by-suppression, with linear responses above spontaneous activity (Fig. S4), whether animals were presented with full-field gratings (drifting at 2 Hz; Methods) or small oriented patches (Gabors; full width at half max, FWHM = 15 degrees; aligned to cortical retinotopic location, also drifting at 2 Hz).

These results were not due to potential nonlinearities in the calcium indicator. We selected jGCaMP8s for this work because it is nearly linear below saturation, and thus for low spike counts its fluorescence represents average spike numbers without distortion. Still, to ensure these effects were not distorted by indicator properties (i.e., nonlinearities between spikes and fluorescence intensity), we deconvolved fluorescence traces using methods that account for indicator nonlinearities. The attenuation seen for suppression and the largely linear behavior for elevated rates (Fig. 2O) remains present in deconvolved data (Fig. S5), including for deconvolution methods that do (Deneux et al., 2016) and do not (Friedrich et al., 2017) account for indicator onset nonlinearities, as expected from the nearly-linear properties of jGCaMP8s for low spike rates. Because deconvolution can be sensitive to estimation convergence and data quality, we further confirmed these results by recreating fluorescence traces using a spike-to-fluorescence model (Zhang et al., 2023) that either did or did not include indicator nonlinearities (Fig. S6). All these analyses support the idea that our measurements are not significantly distorted by the calcium indicator.

In these optogenetic experiments (Fig. 2) we targeted small groups of cells for stimulation (5-12 cells) to increase yield by measuring several neurons in parallel, while avoiding triggering larger network interactions, which have been reported to occur at and above approximately twenty stimulation targets (Marshel et al., 2019; Dalgleish et al., 2020; LaFosse et al., 2023; O’Rawe et al., 2023). Stimulation of small groups like the ones we use has been found to induce smaller network effects, typically weak, broad suppression (Jouhanneau et al., 2018; Chettih and Harvey, 2019; Oldenburg et al., 2024), see (O’Rawe et al., 2023; Sanzeni et al., 2023) for simulations across input pattern size. However, to confirm that our responses are not affected by the number of cells stimulated simultaneously, we performed experiments with only one neuron stimulated at a time (Fig. S7), and found that the results were consistent with our multi-cell stimulation experiments.

Together, these results imply the average IO function of pyramidal cells in mouse V1 exhibits a linear regime above, and a nonlinear regime below, the resting activity level of neurons in the awake state. The nonlinearity below rest attenuates inputs, so that suppression leads to attenuation.

### Consistent effects — attenuation, not amplification — measured in individual pyramidal cells

We next asked if responses reflected by the cell-average trends were present within individual neurons. First, we identified cells whose activity was suppressed by a drifting grating of one direction but increased by the orthogonal direction (Fig. 3A). We then optogenetically stimulated these cells as they fired spontaneously and also while we presented either grating (Fig. 3B-C; N = 6 cells in N = 2 animals). We found results in single cells that are consistent with our previous results from populations of cells: suppression led to attenuation of responses, while optogenetic stimulation applied when cells’ activity was elevated by visual input produced similar responses (i.e., a linear response; p = 0.031; Wilcoxon signed-rank test; Fig. 3D).

We next examined whether stimulation of a single cell with multiple optogenetic intensities (Fig. 3E-F) was consistent with the results we found in averages over cells (Fig. 2). If neurons have a nonlinear region below their resting point, then we should expect a smaller difference between responses (smaller slope) to the two stimulation intensities in suppressed cells. Consistent with this idea, we found that as cells were suppressed, the difference in responses between high and low powers decreased (Fig. 3G, N = 27 cells in N = 3 animals; Fig 3I, left, p = 0.027; Wilcoxon signed-rank test with Bonferroni correction). Similarly, a linear region above cells’ resting point implies there should be little or no difference in responses as cells’ firing rates increase. Confirming this, we found no difference in responses at either optogenetic power (N = 25 cells in N = 3 animals; Fig. 3H) and found there was no difference (change in slope) between high and low powers (Fig. 3I, right; Fig. S8).

In sum, these results confirm that individual neurons show IO functions that are consistent with the population data in Fig. 2. These IO functions are nonlinear below resting activity levels, showing attenuation-by-suppression, and linear for moderate increases in firing rate — across much of the range evoked by visual input.

### Cells operate at the transition point of a supralinear-to-linear IO function at spontaneous activity levels

We next sought to estimate the IO function across the range of activity levels we measured. To do this, we used changes in optogenetic responses (Fig. 2K) to infer the slope of the underlying function. We integrated the curve fitted across all cells’ change in responses (Fig. 2O) to generate the underlying IO function (Fig. 4A-B). Consistent with our individual observations, this IO function has a supralinear region followed by a linear region, before eventual saturation at high inputs (Fig. 4B).

One constraint with our approach is that we obtain the IO function by integrating changes in optogenetic response as a function of changes in ongoing neural activity. This method characterizes changes in response, but does not define zero firing rate (x-axis, Fig. 4B). To ensure that the shape of the IO function we found is valid even at low activity levels, we characterized the impact of interpolated extensions to the fitted line on the resulting IO function. We found no qualitative differences (Fig. S9) from our prior estimates. To estimate the zero point more directly, we estimated neurons’ firing rate based on their calcium transient rate by deconvolving each neuron’s response. This analysis revealed that there are indeed cells whose activity reaches zero, approximated by a zero rate of calcium transients, during visual stimulation (Fig. S9). We used this value to represent zero firing rates in Fig. 4B, and also found that deconvolved responses yield the same shape of the underlying IO function as when using calcium responses (Fig. S10).

The IO function shape we found is comparable to the Ricciardi function, an IO function that arises in spiking network models that are recurrently connected. Its shape is influenced by the inputs neurons receive from other neurons in the network, because variability in membrane potential and membrane conductance influence IO functions (Chance et al., 2002), and these vary with input. The Riccardi function shows a supralinear, then a linear, regime with large or small amounts of saturation depending on input and other parameters such as maximal firing rate and refractory periods (Sanzeni et al., 2020). We fit the parameters of the Ricciardi function to our data (Methods), finding a strong similarity between the model and our data (Fig. 4B). Compared to other models of the IO function, like threshold-linear or power law functions, the Ricciardi function produces the closest fit to our data (Fig. S11).

To assess the variability of the IO function fit across neurons, we calculated bootstrapped confidence intervals of the Ricciardi function fit (Fig. 4B; on each bootstrap trial, fitting a LOESS, integrating, and fitting the Ricciardi function to the resulting IO function estimate; parameter sensitivity analysis in Fig. S12).

Our results are summarized in Fig. 4C. Neurons operate during spontaneous activity near the transition from supralinear to linear regions of their IO function. Suppression then moves neurons into the supralinear region, so that the more a neuron is suppressed, the smaller its response to a fixed input. Thus, the nonlinearity in the IO function, and the operating point of the neurons during spontaneous activity, together produce attenuation-by-suppression.

### Trial-to-trial response variability reveals differences in IO function gain, but not IO function shape

The supralinear-to-linear IO function shape we find is an average across observations from many neurons. To explore variability that might exist across cells, we leveraged the trial-to-trial fluctuations in cells’ activity to examine responses at multiple points along a cell’s IO function (Fig. 5A). From one trial to the next, neurons’ activity (ΔF/F_0_) fluctuates. Therefore, on different trials, cells may be more or less excited (or suppressed) just before the onset of optogenetic stimulation. By relating the magnitude of neurons’ optogenetic responses to the prior ΔF/F_0_ levels, we obtain multiple measurements along the IO function for each cell, allowing us to examine the shapes of individual cells’ IO functions.

**Fig. 5:**
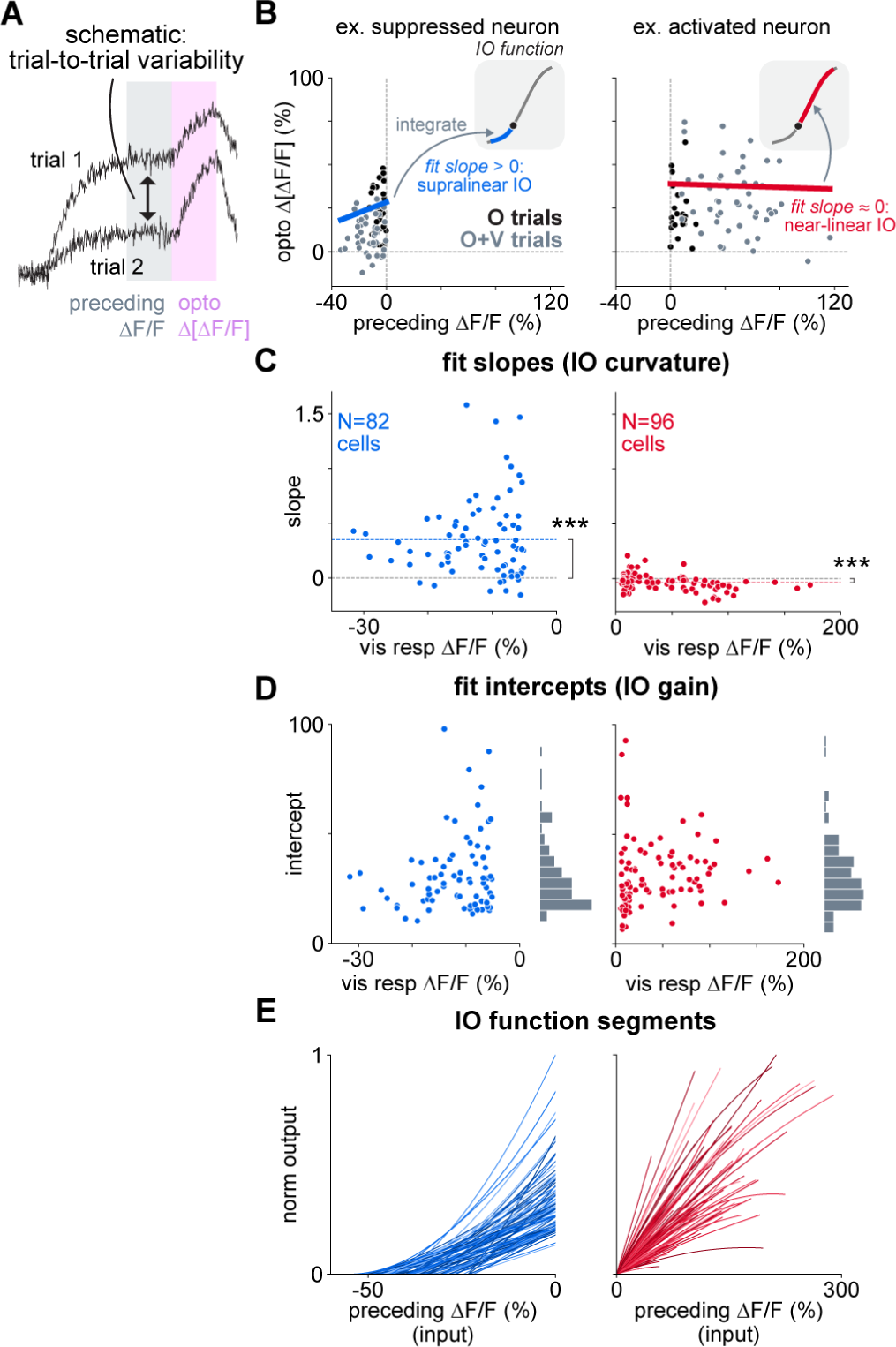
Trial-to-trial variability reveals that IO function shape is consistent across cells, while IO function gain is variable. **(A)** Schematic of analysis: there is trial-to-trial variability prior to optogenetic stimulation that causes each trial’s optogenetic stimulus to arrive at a different baseline level of activity. **(B)** Example cells. X-axis: activity prior to optogenetic stimulation. Y-axis: optogenetic response. (Left) suppressed neuron, (right) activated neuron. Colored lines: individual fits across trial data. Insets: schematic of the IO function segment that each fit line represents, computed by integrating the fit. **(C)** Fit slopes across all suppressed (left, blue: visual response < -5% ΔF/F0; mean ± SEM = 0.35 ± 0.04; N = 82) and activated (right, red: visual response > 5% ΔF/F0; mean ± SEM = -0.04 ± 0.01; N = 96; right) cells. This parameter gives the curvature of cells’ IO functions. Dashed colored lines: mean slope. ***p < 0.001, Wilcoxon signed-rank test with Bonferroni correction. **(D)** Same as C, but for fit intercepts across cells, which give the gain or slope of cells’ IO functions. Gray histograms: variability of fit intercepts across cells. (E) IO function segments derived by integrating fits in B; all lines normalized to the single largest peak response.

To characterize cells’ IO function shapes, we first estimated the change in optogenetic response as a function of preceding activity. We used a mixed-effects model,

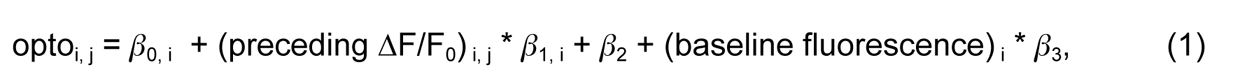

to estimate effects of preceding activity while accounting for cells’ baseline fluorescence levels (β_0_, β_1_ random effects, other coefficients fixed effects, i indexes cells, j trials; Methods). For cells transfected with the bicistronic virus that express both opsin and indicator, the baseline fluorescence is a proxy for opsin level (same cells as in Fig. 2N, data from bicistronic virus expression animals only, N = 6 animals, N = 9 experiments; N = 178 total cells). For each cell we used only trials where the activity before optogenetic stimulation was either below (suppressed; range 55-81 trials) or above (activated; 44-82 trials) spontaneous activity, to limit distortion of fit slopes due to activity switching between a supralinear and linear regime (Fig. 3A-D).

This model yields a fit line for each cell (Fig. 5B). This fitted line represents the derivative of the cell’s IO function, so that the slope of the line indicates the curvature of the IO function. That is, a slope near 0 reflects linearity, a slope > 0 reflects supralinearity, and a slope < 0 reflects sublinearity. The intercept of the fitted line corresponds to the slope, or the gain, of the IO function at cells’ resting point (i.e. where the preceding ΔF/F_0_ = 0%, or, where there is no change from baseline activity).

Examining the distribution of fit slopes across all visually-activated and suppressed cells reveals IO function shapes consistent with our previous findings. First, we found visually-suppressed cells primarily operate along a supralinearity in the IO function (Fig. 5C, left; N = 82 cells; mean fit slope = 0.35 ± 0.04; p = 4*10^-13^, Wilcoxon signed-rank test with Bonferroni correction). On the other hand, cells whose activity was elevated above resting levels show near-linear IO functions, though many operate along slightly sublinear IO functions, suggesting a transition from linear to saturating responses (Fig. 5C, right; N = 96 cells; mean fit slope = -0.04 ± 0.01; p = 7*10^-7^, Wilcoxon signed-rank test with Bonferroni correction).

Next, we used the distributions of fit intercepts to determine if gain, or IO function slope, varies across cells (Fig. 5D). This is a new measurement, enabled by the use of trial-to-trial variation, as the analyses in Fig. 2 average across cells when computing slopes. The principal result is that while excited cells are all quite linear, their gain (slope) varies from cell to cell. Note that this variation in gain could be due to differences in opsin expression, or due to underlying variation in cells’ gain, perhaps due to receiving different levels of network input (Chance et al., 2002). For both suppressed and activated cells, we found significant variance (p < 0.001 for both groups, via permutation tests; Fig. S13). We integrated cells’ fits to construct the segment of their IO function corresponding to the fits (Fig. 5E; qualitative comparisons of IO function shape across neurons in Fig. S13). Remarkably, cells are quite consistent in shape within groups. Suppressed cells show qualitatively similar supralinear shapes. Activated cells are linear or near-linear, with moderate saturation at high levels of activity, but their gain varies significantly (Fig. 5D).

In sum, we find that above rest or spontaneous activity levels, cells consistently operate in a near-linear regime with some saturation, and below rest, show a supralinear IO function shape. Despite the variability we measured in the overall slope, or gain, of the IO functions, the attenuation-by-suppression effects are consistent across cells.

## Discussion

Here, we demonstrate through direct stimulation that excitatory cells in the cortex of awake mice operate along a supralinear-to-linear IO function. Cells’ f-I curves, or IO functions, have previously been suggested to exhibit smooth nonlinearities, such as power-law nonlinearities, across the working range of visual responses (Anderson et al., 2000; Carandini, 2004; Finn et al., 2007), and it was thought these nonlinearities would serve to amplify inputs (Ahmadian et al., 2013; Rubin et al., 2015). However, we find a key feature of V1 excitatory cells is not that they exhibit amplification above baseline (spontaneous) activity levels, but rather, that they show linear responses. We do find that neurons’ IO functions show a smooth nonlinearity — a supralinearity — but this effect occurs below the level of spontaneous firing on average. Thus, we conclude that the prominent nonlinearity in neurons’ IO functions does not allow amplification, but instead causes neurons to attenuate their responses to input as they are suppressed.

Our inferred IO function shape (Fig. 4B) has a supralinear region, which we show here is below resting or spontaneous activity, a linear region that covers most of neurons’ operating range as defined by high visual stimulus contrast (100% contrast, Fig. 2, Fig. 3A-D), and a weakly sublinear region for the highest activity levels.

### Relationship to prior work

Here we measure the somatic IO function, or how neurons’ spike rates change as a function of aggregate inputs from the dendrites. While dendritic nonlinearities may change how certain inputs are integrated, the somatic IO function is a good description of neurons’ full IO function when inputs are summed linearly in the dendrites. While nearly linear summation in dendrites can occur in some conditions (Katz et al., 2009; Menon et al., 2013), nonlinear processes in the dendrites can add to the neural computations performed by recurrent networks of neurons (e.g., (Gütig and Sompolinsky, 2006; Smith et al., 2013; Kerlin et al., 2019)). Understanding brain function will require understanding both the recurrent network computations that occur via near-linear dendritic summation as well as the computations created by dendritic nonlinearities.

Two prior observations lend credence to our findings. First, our data are strikingly similar to the Ricciardi IO function that is derived from integrate-and-fire model networks (Ricciardi and Smith, 1977; Sanzeni et al., 2020). Ricciardi functions can have dramatic saturation at high activity levels, and thus a major result of our work is that the normal operating range of cortical neurons is largely below saturation, in a near-linear regime for elevated rates. Previously, it was unknown if the Ricciardi function was a good description of cortical neurons’ responses *in vivo*. Second, a set of dynamic clamp experiments which simulated conductance inputs finds IO function shapes that are also similar to our results (Gajowa, 2018).

Prior studies have estimated a power law relationship between membrane potential and spike rate *in vivo,* with an expansive supralinearity throughout neurons’ input range (Anderson et al., 2000; Finn et al., 2007; Tan et al., 2014). While the shape of a power law in some regimes can approximate the supralinear-to-linear IO function we see, prior characterizations of a power law in the relationship between membrane potential and spike rate do not necessarily mean the IO function follows a power law. This is especially true at higher spike rates, when neural firing rates become more dependent on refractory periods and time to integrate to threshold (Sanzeni et al., 2020), instead of spikes being governed by membrane potential fluctuations (Dayan and Abbott, 2005). Also, direct input injection, as we use, is better able to determine where the supralinearity in the IO function is relative to spontaneous activity. This allowed us to determine that neurons sit near this transition during spontaneous firing. In addition, our all-optical approach has allowed us to measure the responses of a large population of neurons, and these data suggest that while individual neurons share the same supralinear-to-linear IO shape, there is variability in gain or slope across cells (Figs. 4-5). Such variability across cells may serve to improve information coding at the population level, as has been suggested of the variability found in neurons’ tuning curves (Ecker et al., 2011).

We observed some saturation in IO functions for neurons that showed the largest responses to strong input. This effect could be caused by mechanisms such as sodium channel inactivation in single cells, strong inhibitory coupling in the network, and/or consequences of the refractory period (Rauch et al., 2003; Sanzeni et al., 2020). However, we cannot rule out that decreased responses were due to saturation of fluorescent signals at high activity levels (Zhang et al., 2023).

### Recurrent network input likely plays a role in creating the supralinearity

What shapes the supralinearity we measure for low inputs? In principle, this soft threshold for the somatic IO function could arise from biophysical factors like voltage-gated channels. If that were the case, we might expect to see the supralinearity in *in vitro* measurements of f-I curves, but these often show a threshold-linear response function, not a soft threshold or supralinearity (McCormick et al., 1985). Because *in vitro* networks in slices or culture are often in a lower-activity state than the network is *in vivo*, these observations suggest the supralinearity we observe may arise from inputs to cells from other neurons in the network. Neurons in the cortex receive a constant barrage of excitatory and inhibitory inputs, which can lead to ongoing fluctuations in their membrane potentials (van Vreeswijk and Sompolinsky, 1996; Destexhe and Paré, 1999; Brunel, 2000; Destexhe et al., 2003). These fluctuations can smooth the relationship between input and output near the IO function threshold (Hansel and van Vreeswijk, 2002; Miller and Troyer, 2002). In fact, the Ricciardi IO function that matches our data is an analytical description of the IO function shape produced from inputs in a recurrent network of excitatory and inhibitory neurons (Ricciardi and Smith, 1977; Sanzeni et al., 2020).

### Implications for network computation and brain function

Two major consequences for brain function arise from these results. First, the attenuation-by-suppression we observe can have direct consequences on cortical computation — in other words, the brain can filter out inputs to neurons in certain contexts, by suppressing the activity of those neurons so their incremental responses to other inputs are attenuated. Second, and more speculatively, because the supralinear-to-linear somatic IO function we characterized shares general features with activation functions used in modern machine learning systems, including generative AI systems like ChatGPT (Brown et al., 2020), our results may have implications for learning.

Attenuation-by-suppression has computational consequences for brain operation. Neurons are suppressed in a wide variety of sensory contexts. A classic example is the case of sensory surround suppression. In surround suppression (Hubel and Wiesel, 1965; Allman et al., 1985; Cavanaugh et al., 2002), sensory stimuli outside a neurons’ key feature tuning (outside its receptive field) decrease that neuron’s firing. Surround suppression, combined with the attenuation-by-suppression effect we observe, could lead to sharpening of responses to small visual stimuli or stimuli that change quickly in space, like edges. In this scenario, as some cells are activated by the stimulus, those activating inputs, falling on neurons’ linear region, would be passed on to downstream areas. In contrast, other nearby neurons will have receptive fields that are in the surround — the small stimulus would fall outside their receptive field, causing them to be suppressed (Ozeki et al., 2009; Haider et al., 2010, 2013; Angelucci et al., 2017; Lakunina et al., 2020). These suppressed neurons would then attenuate any additional inputs that arrive, filtering out those inputs, so that surround inputs have less impact on downstream neurons. This effect could, at the population level, serve to amplify relevant responses above background noise (Ringach and Malone, 2007).

Notably, this mechanism is not just the usual conception of suppressive surrounds. Here we show not just that visual stimuli suppress some neurons’ activity, but that suppressed neurons are actively *filtering* out the inputs they receive — that is, attenuating added inputs, or attenuation-by-suppression. We arrive at this result by going beyond the measurements possible with visual stimuli alone, using an *in vivo* cell-specific stimulation approach that allows us to directly measure IO functions and changes in response to fixed input.

Attenuation-by-suppression is related to the idea that expansive nonlinearities due to spike threshold can sharpen responses (Mechler and Ringach, 2002; Priebe et al., 2004); this type of sharpening due to the IO function may be more common across cortical areas than has been previously reported. Attenuation-by-suppression may also offer an explanation for perceptual masking effects arising from optogenetic stimulation (Chen et al., 2022), as optogenetic stimulation can lead to suppression caused by recurrent excitatory connections (Dalgleish et al., 2020; O’Rawe et al., 2023). These effects, where optogenetic stimulation generates suppression and thus attenuation, may also underlie the differences in visual perception noted when visual and optogenetic inputs are combined (Lafer-Sousa et al., 2022; Azadi et al., 2023).

Another major result of this work is that our measured IO functions share important features with the activation functions used in recent AI systems. Over a decade ago, activation functions with a linear regime over much of the unit’s activity range (rectified-linear units, ReLUs) were shown to yield some learning improvements compared to sigmoidal functions. Since then, it has been observed that learning performance can be additionally improved by activation functions that smooth the ReLU’s sharp transition from zero activation to the linear region (Clevert et al., 2015; Hendrycks and Gimpel, 2016). Intuitively, this smoothing avoids a zero gradient below the rectification threshold and limits the degree to which synapses drop out of learning due to a lack of available error signal. One might map this to biology by saying that neurons that “cannot fire never wire.” Thus, while learning rules *in vivo* are not fully understood, if there were a sharp IO function threshold *in vivo*, it might be that learning would be difficult. This smooth nonlinearity followed by a linear region is a shape shared by many activation functions in modern AI systems, including the softplus, Gaussian error linear (GELU), and exponential linear (ELU) functions. Our data show striking similarities to these activation functions, with a smooth transition around the nonlinearity and a largely linear regime through most of the range used for even high-contrast visual responses. Since these activation functions work well in model networks, we might speculate that biological neurons have faced selection pressure to produce IO functions with this shape, to optimize some form of learning or other computation.

## Conclusion

In summary, our findings characterize the average IO function of neurons in the cortex during normal brain operation — in the awake state, as animals actively receive sensory input. The attenuation-by-suppression we observe suggests a new role for ongoing activity in cortical networks. Because ongoing input creates the smooth nonlinearity below spontaneous firing rates, it can perform computations by allowing neurons to selectively filter out extraneous features of sensory input.

## Methods

### Viral injections and cranial window implants

All procedures were approved by the NIMH Institutional Animal Care and Use Committee (IACUC) and conform to relevant regulatory standards. Emx1-Cre mice (The Jackson Laboratory) were used in all experiments to target expression of Cre to excitatory neurons. Adult mice were anesthetized with isoflurane (1–3% in 100% O2 at 1 L/min) and administered intraperitoneal dexamethasone (3.2 mg/kg). A custom metal head post was fixated at the base of the skull. A 3 mm diameter circular craniotomy was made over the left hemisphere of primary visual cortex (ML -3.1 mm, AP +1.5 mm relative to Lambda).

A bicistronic virus expressing both opsin and GCaMP indicator, AAV9-hSyn-DIO-jGCaMP8s-P2A-stChrimsonR(LaFosse et al., 2023) (Addgene, 174007) was diluted in phosphate-buffered saline (final titer: 2.6*10^12^ GC/mL, N = 3 mice; 2.9*10^12^ GC/mL, N = 3 mouse; 3.4*10^12^ GC/mL, N = 2 mice; 4.7*10^12^ GC/mL, N = 2 mice; 6.5*10^12^ GC/mL, N = 2 mice) and injected 200 µm below the surface of the brain (multiple injections, 300 nL/injection, 0.1 µL/min) to target expression to layer 2/3 neurons. In N = 1 mouse, separate viruses ofAAV9-hSyn-jGCaMP8s-WPRE (1.0*10^13^ GC/mL) and AAV9-hSyn-DIO-stChrimsonR-mRuby2 (2.7*10^12^ GC/mL) were instead injected.

A 3 mm optical window (Tower Optical) was implanted over the craniotomy. Both the optical window and metal head post were fixed to the skull using C&B metabond (Parkell) cement dyed black. Animals were individually housed after surgery. Mice were imaged three or more weeks post-injection, and water-scheduled (1.0 mL/day) to promote alertness during experiments that used intermittent water delivery.

### Two-photon calcium imaging

Two-photon calcium imaging was performed with a custom-built microscope (MIMMS, Modular In vivo Multiphoton Microscopy System, components, Sutter Instruments; controlled by ScanImage, MBF Biosciences; bidirectional 8 kHz resonant scanning, 512 lines, ∼30 Hz frame rate). Calcium responses were imaged with 920 nm laser pulses (Chameleon Discovery NX laser, Coherent, Inc., pulse rate 80 MHz, pulse energy 0.19–0.25 nJ/pulse; imaging power 15–30 mW measured at the front aperture of the objective; 16x objective, 0.8 NA, Nikon Inc; immersion with clear ultrasound gel, ∼1 mL; 100-200 µm below pia, L2/3 of V1, field of view 400 x 400 µm). In principle, optical crosstalk leading to inadvertent activation of opsin during imaging might affect our measurements. However, we chose the stChrimsonR opsin to minimize such crosstalk activation, and under similar imaging conditions as in this study and using the bicistronic virus we used here (N = 9 of 11 experiments), we find no evidence for such crosstalk activation (LaFosse et al., 2023).

### Cell-specific two-photon stimulation

Cell-specific photostimulation was performed using either 1030 nm (Satsuma laser, Amplitude Laser; Fig. 2, Fig. 3E-I) or 1035 nm light (Monaco laser, Coherent, Inc.; Fig. 3A-D). The laser wavefront was shaped by a spatial light modulator (1920 x 1152 pixels; Meadowlark) to target patterns of cells for stimulation (10 µm diameter disks; 300 ms, 8 mW/target, 6.5 mW/target for low power in Fig. 3E-I, 500 kHz pulse rate; TTL logic gate, Pulse Research Lab, used to invert imaging line clock to gate laser during pixel acquisition and stimulate during reversal of bidirectional scanning: on time 19 µs, off time 44 µs, 30% duty cycle; stimulation power accounting for blanking: 2.4 mW/target, or 2 mW/target for low power; see LaFosse et al., 2023 for more details). We see little off-target activation in the X-Y plane with the 10 µm disks we use here (LaFosse et al., 2023). It is possible that the larger axial extent of the stimulation spot may at times recruit off-target neurons above or below our stimulation disks. The fact that stimulating with one target at a time leaves our results unchanged (Fig. S7) suggests that any off-target activation has little effect on our results.

### Visual stimulation

Visual responses were measured in awake, head-fixed mice viewing sinusoidal drifting grating stimuli (full-field or Gabor patches filtered with a 15° full-width half-max Gaussian mask; spatial frequency of 0.1 cycles/degree; speed 20 deg per sec; drifting at 2 Hz; 100% contrast; 3-4 seconds; stimuli presented on an LCD monitor with neutral gray background).

To align Gabor patches to the receptive fields of neurons in the imaging FOV, we performed retinotopy via hemodynamic intrinsic imaging using 530 nm light (delivered via fiber-coupled LED, Thorlabs; 2 Hz imaging) (Goldbach et al., 2021). Retinotopic maps were calculated by measuring hemodynamic-related changes in light absorption across the cortex to square wave drifting gratings (0.1 cycles/degree, 10° diameter, 6 locations, 5 seconds; 5 second baseline window and 2.5 second response window starting 3 seconds after stimulus onset used to measure responses).

### Experimental sequence

To ensure mice were awake and alert, small volumes of water were delivered (3 µL per trial, ∼300 µL total per session; 20% of trials with independent probability; 50-60 seconds average time between delivery; water delivered 2.5-3.5 seconds before onset of optogenetic stimulus and 2 seconds before onset of visual stimulus).

For all experiments, visual responses were measured to drifting grating stimuli of different directions (8 directions 45° apart randomly interleaved, 20 repetitions per direction, 2 seconds on, 6 seconds off). Mice were head-fixed and sat in a tube during experiments. For optogenetic stimulation experiments, a single direction was chosen that led to activation or suppression of a number of neurons in the field of view (2-3 patterns of neurons per experiment; Fig. 2 and Fig. 3E-I). During these experiments, trials with visual only, optogenetic only, and paired stimulation trials were randomly interleaved; visual stimulus always the selected single drifting grating; 50 repetitions per trial type per optogenetic stimulation pattern, 250-350 trials total; 6-9 second inter-stimulus interval for optogenetic stimulation. Experiments lasted 100-120 min total (20 min visual stimulation of multiple directions, 30 minutes for visual response characterization and neuron selection, 50-70 minutes for optogenetic and visual stimulation experiments).

For experiments in Fig. 3A-D, two orthogonal visual stimulus directions were chosen. Experimental sequence was the same. We collected visual response data as described above, multiplied average pixel-wise responses for all orthogonal pairs of directions, and selected neurons for stimulation whose value after multiplication was negative, indicating a positive response for one direction (visually-activated) and negative response for the orthogonal direction (visually-suppressed).

### Analysis of two-photon imaging data

Images acquired via two-photon imaging were first rescaled from 512 x 512 pixels to 256 x 256 to ease data processing. Background correction was done by subtracting the minimum pixel value of the average intensity image and setting all remaining negative pixels (due to noise) to zero. We used the CaImAn toolbox (Giovannucci et al., 2019) for motion correction and Suite2p (Pachitariu et al., 2017) for cell segmentation to allow manual selection of cell masks.

We measured neurons’ activity to both visual and optogenetic stimuli (ΔF/F_0_; F: average intensity across all pixels within a cell’s segmented mask; F_0_: average fluorescence across the 45 imaging timepoints, 1.5 s, prior to stimulus presentation). “Opto” trials or “Vis” trials are where the corresponding stimulus was delivered by itself. “Opto+Vis” trials, (O+V), are when visual stimulus and optogenetic stimulus are delivered on the same trial. The “Opto during vis” response, (O+V) - V, Fig. 2M was found by subtracting the average Vis trace from the average Opto+Vis trace. Deconvolved activity was estimated using the constrained OASIS method (Friedrich et al., 2017) or MLspike (tau = 0.4 s; amplitude parameter for each cell defined as smallest peak of the denoised calcium trace from the OASIS method; (Deneux et al., 2016)).

For response ΔF/F_0_ maps (Fig. 2D, F-K; Fig. 3B), we measured trial-averaged activity at every pixel (ΔF/F_0_; F: average frame across a time window, Fig. 2D, entire visual stimulus period, Fig. 2F-I and Fig. 3B, entire visual stimulus period except the first 1 s, to exclude the initial transient response at stimulus onset, Fig. 2G-J, 1 second period following optogenetic stimulus onset; F_0_: average frame from 1.5 s before onset of the stimulus).

### Analysis of stimulus-evoked responses

Cell responses (points in Fig. 2N-O; Fig. 3C-D, G-I) were computed by averaging timecourses in 0.33 second windows (10 frames), just before (baseline) or after (response) optogenetic stimulus onset. “Vis resp” (x-axis, Fig. 2O, 3D, 4A): response on Opto+Vis trials, in window just before optogenetic stimulus onset (when only the visual stimulus was present). “Opto” (Fig. 2N): response on Opto only trials, in response period after optogenetic stimulus onset, minus baseline period just before optogenetic stimulus onset. “Opto during vis”, “(O+V) - V” (Fig. 2N): response period on Opto+Vis trials minus response period on Vis only trials, with baseline period difference on the same trials subtracted. Change in optogenetic response between visual and spontaneous conditions (y-axis, Fig. 2O, 3D, 4A): difference between “Opto during vis” and the “Opto” responses.

Stimulated cells during our experiments were defined as those whose centroid fell within a 15 micron radius of any stimulation target’s center (5-12 targets per pattern, average 9 ± 0.5) and whose “Opto” response value exceeded a 5% ΔF/F_0_ threshold (N = 15 ± 1.3 stimulated cells per pattern).

### IO function estimation

To estimate the underlying IO function of V1 pyramidal cells, output = ƒ(input), we first performed locally estimated scatterplot smoothing, or LOESS (Cleveland, 1979). We fit LOESS curves (rotated axis to maximize variance on x-axis, using the loess Python package (Cappellari et al., 2013): pypi.org/project/loess/) to the data (Fig. 2O, Fig. 4A; confidence intervals via bootstrap; N = 1000 repetitions, N = 364 samples).

The resulting fit represents a change in output versus the input level of a cell, or the derivative of the IO function, shifted vertically by a constant:

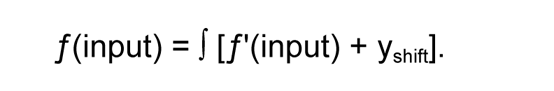

The value of y_shift_ is dependent on two factors (Fig. S9A): any difference between maximum visual suppression and zero firing rate (y_supp,0_), and the size of the optogenetic response (y_opto_): y_shift_ = y_supp,0_ + y_opto_. Deconvolved spiking estimates of calcium activity (Fig. S9B-E) suggest the most-suppressed cells did indeed reach zero response, so we estimate the shift as the minimum of the fit line: y_supp,0_ = -*min*(ƒ’(input)). Assuming suppression to zero completely attenuates small optogenetic inputs (as expected for an activation function that goes to zero), we set y_opto_ to zero. While we cannot directly measure this value because any optogenetic activation moves responses away from zero, we confirm that our conclusions are not changed by various extrapolations from our observed data points (Fig. S9F-H).

To fit the Ricciardi transfer function to our estimated IO function (Fig. 4B), we used least squares regression. Initial values for the parameters taken from Sanzeni et al. (2020): τ_i_ = 20 ms, τ_rp_ = 2 ms, *V_r_* = 10 mV, θ = 20 mV, σ = 10 mV. The fitted Ricciardi function in Fig. 4B and 95% confidence intervals were generated by taking the mean of the estimated IO functions computed at each bootstrap repetition of the LOESS fits on the data.

For comparison of the Ricciardi transfer function fit with other model IO functions (Fig. S11), we fit the following equations to our data, where the parameters a, b, and c were fit for each function:

Threshold-linear (rectified-linear, or ReLU):

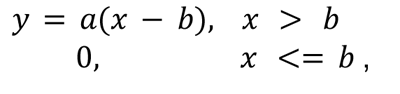

Power law:

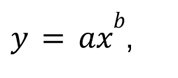

Logistic:

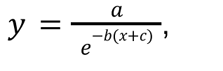

Stretched exponential:

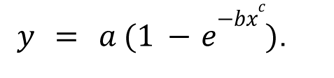

### Ansalysis of IO variability

In order to generate fits across trial-to-trial data for individual cells, we fit a mixed-effects model with a fixed effect for baseline fluorescence (a proxy for opsin levels in neurons expressing the bicistronic construct, LaFosse et al., 2023), to control for differences in optogenetic response due to differences in opsin expression. We thus only included cell data from animals expressing the bicistronic construct in this variability analysis. Cell variation in slope and intercept were random effects.

Example fits, as in Fig. 5B, are computed using all the terms in Equation 1. Analysis of variability across cells (Fig. 5C,D; Fig. S13) uses cells’ slopes β_1, i_, and intercepts for each cell, β_0, i_ + β_2_ + (mean baseline fluorescence) * β_3_.

## Data, Materials, and Software Availability

Raw data of cell traces and code to generate figure panels will be deposited and made publicly available on Zenodo at the time of publication.

## Author contributions

PKL: experiments and data analysis. JO: data analysis. ZZ, NF, VS: experiments. YD, MH: two-photon holographic stimulation equipment. PKL, MH: manuscript writing, with feedback from all authors.

## Acknowledgements

We thank Mark Stopfer, Bruno Averbeck, Bei Xiao, and Chris Kim for comments and discussion. We thank Dylan Nielson for comments and feedback on our mixed effects model. We thank Georg Jaindl and others from Vidrio/MBF Biosciences for expert technical assistance. This work used the computational resources of the NIH HPC Biowulf cluster (http://hpc.nih.gov). This work was supported by the NIH BRAIN Initiative grant U19NS107464 and the NIMH Intramural Research Program, ZIAMH0020967.

## Supplemental Figures

**Fig. S1:**
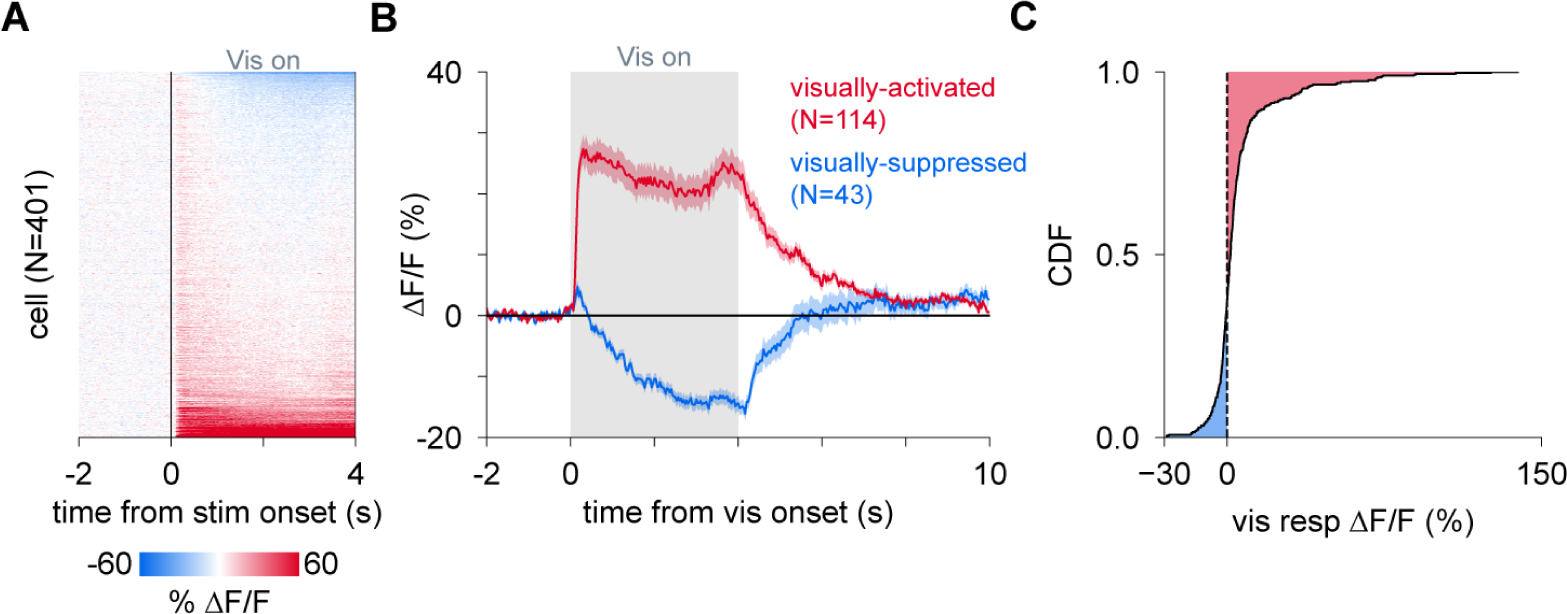
Drifting gratings evoke a pattern of visually-activated and visually-suppressed cells. **(A)** Example visual responses in one animal (see Fig. 2D) to a drifting grating stimulus. **(B)** Cell-averaged (mean ± SEM) activity traces to the visual stimulus for visually-activated (red) and visually-suppressed cells (blue). Cells were determined as activated or suppressed by averaging the activity across the visual stimulation period (excluding the first ∼330 ms to account for initial transient response). Activated: > 5% ΔF/F_0_, suppressed: < -5% ΔF/F_0_. **(C)** Cumulative distribution function of visual responses across all cells.

**Fig. S2:**
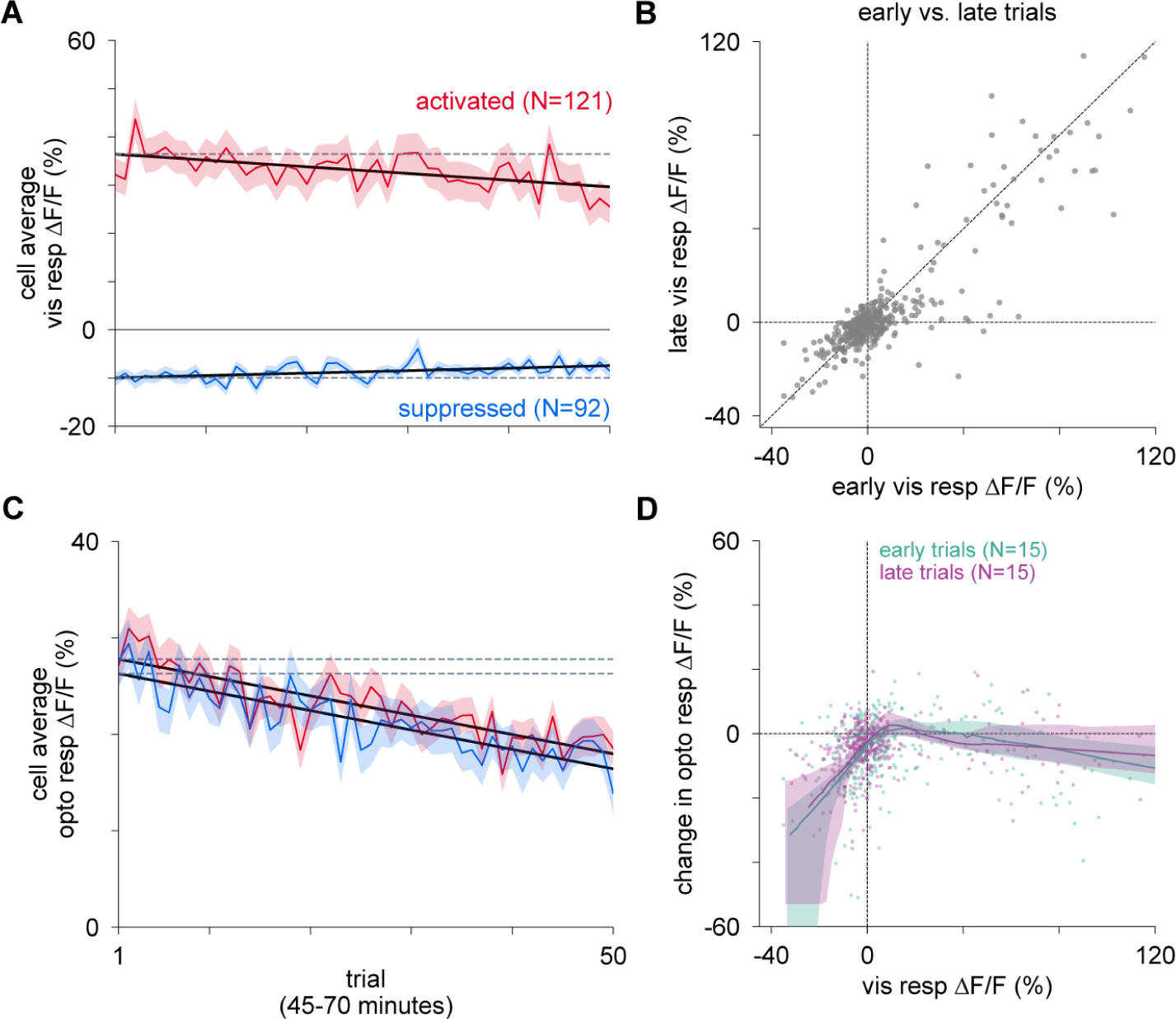
IO functions are stable within experimental sessions. **(A)** Cell-averaged (mean ± SEM) visual responses across trials. There is minimal response adaptation, or change over trials, for both visually-activated (red; linear fit (black line) R^2^ = 0.32; slope nonzero, p = 2*10^-5^) and visually-suppressed (blue; linear fit R^2^ = 0.21, p = 7*10^-4^) cells. Grey dashed horizontal lines: value of the fitted response on the first trial. **(B)** Visual responses (N = 364 cells; Fig. 2N) averaged across early (first 15 trials) and late (last 15 trials) periods. Diagonal line with slope 1 is shown for visual purposes; statistics on early vs late changes described for panels A and C. **(C)** Cell-averaged (mean ± SEM) optogenetic responses across trials showing a slow decline in evoked responses in both activated (red; linear fit R^2^ = 0.70, p = 3*10^-14^) and suppressed (blue, linear fit R^2^ = 0.71, p = 1*10^-14^) cells. Grey dashed horizontal lines: same as in **A**. **(D)** Despite some changes in visual and optogenetic responses within sessions, IO function estimates are not changed. Scatter plot of change in optogenetic response versus visual response for all cells shows the same attenuation-by-suppression for early versus late trials.

**Fig. S3:**
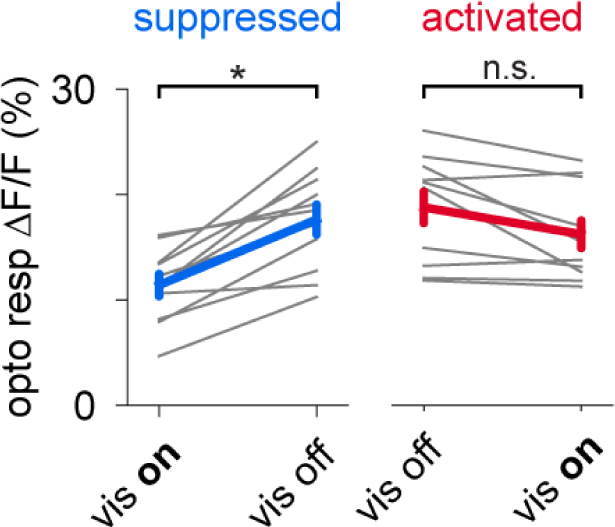
Attenuation of responses during visual suppression is seen across subjects. Cell-averaged optogenetic responses during and without visual stimulation for visually-suppressed and visually-activated cells for each experiment. Gray lines: individual experiment data from N = 11 experiments from N = 7 animals. Blue and red lines: mean ± SEM across all experiments showing decreases in optogenetic responses in suppressed cells during visual stimulation (*p < 0.05; student’s t-test with Bonferroni correction for multiple comparisons) but no changes in activated cells.

**Fig. S4:**
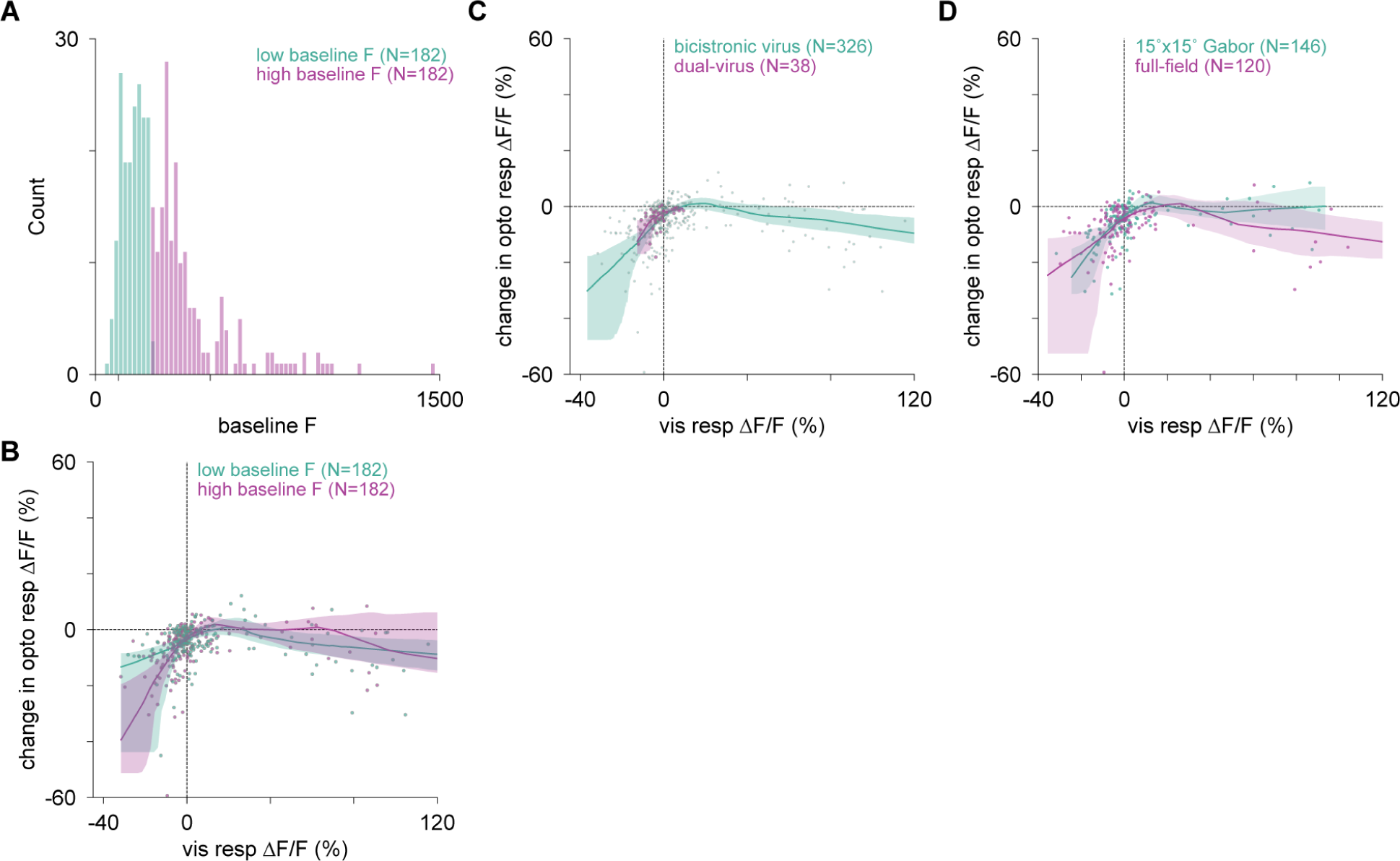
Changes in optogenetic responses due to visual stimulation are not impacted by viral expression or the size of visual stimuli. **(A)** Histogram of all cells’ baseline fluorescence values, divided into two groups: cells with lower baseline fluorescence and higher baseline fluorescence. This may indicate differences in spontaneous firing rates or differences in overall expression of the viral construct. **(B)** Change in optogenetic response as a function of visual response for the two groups of cells with lower (teal) or higher (purple) baseline fluorescence. **(C)** Change in optogenetic response as a function of visual response with cells divided into groups by viral expression strategy: bicistronic virus in N = 9 experiments (teal) and a dual-virus approach (see Methods) in N = 2 experiments (purple). Both the nonlinearity due to suppression and the near-linear effects for elevated firing are seen in both types of preparations, though the data are more limited for the dual-virus approach. **(D)** Change in optogenetic response as a function of visual response across cells, measured with both full-field drifting gratings (purple) and 15 x 15° Gabor drifting gratings (teal). Lines: LOESS fits with 95% confidence intervals. No qualitative differences in results are seen.

**Fig. S5:**
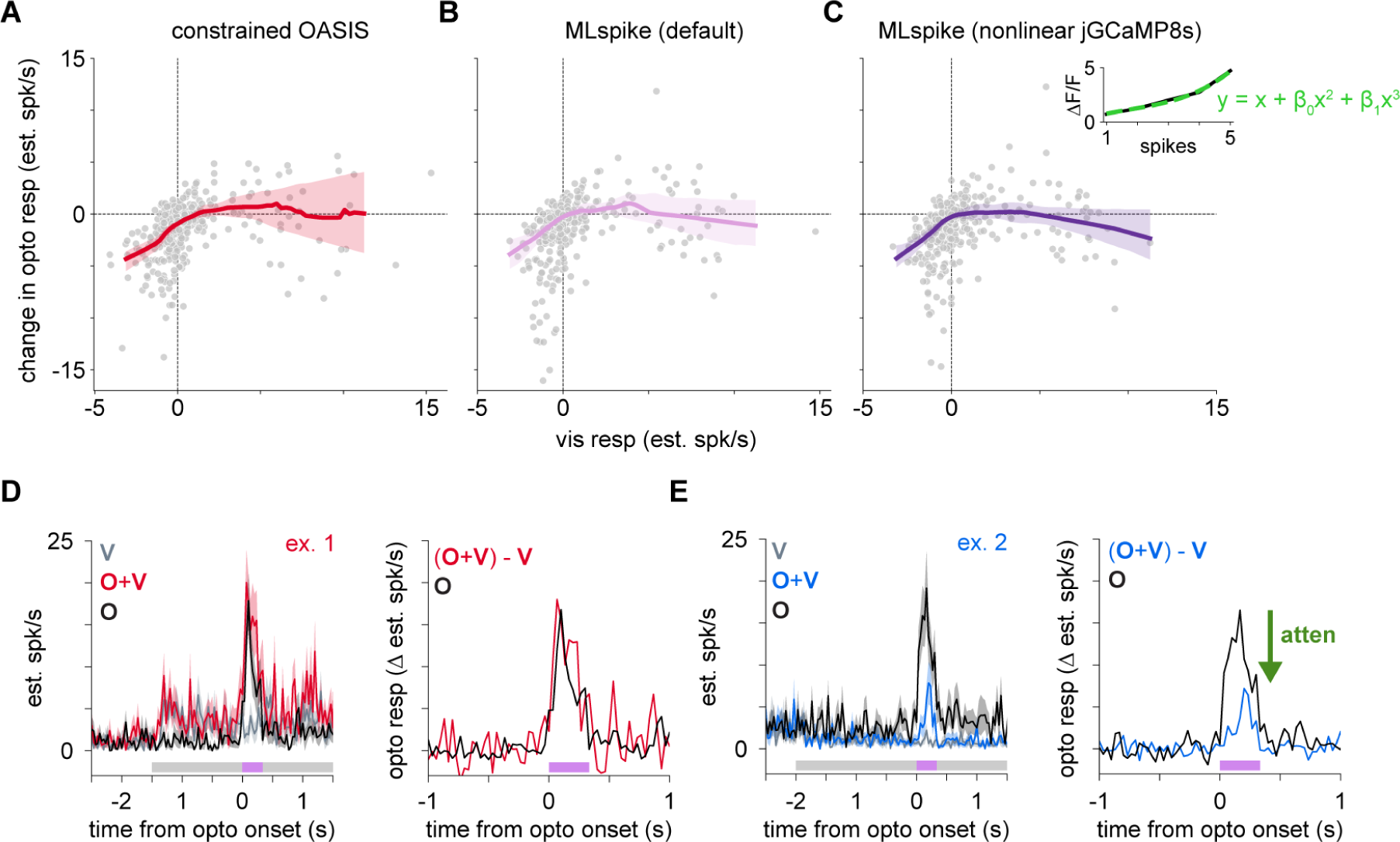
Deconvolution with both linear and nonlinear methods reproduces measures of attenuation during visual suppression. **(A-C)** Three methods for deconvolution yield IO functions that are similar to those estimated with ΔF/F_0_ measurements (e.g. Fig. 2O). **(A)** A linear deconvolution method (constrained OASIS; Friedrich et al., 2017) was used to reconstruct spike rate estimates on each trial and those responses used to calculate IO functions. Red line: LOESS fit to data with 95% confidence intervals via bootstrap. **(B-C)** To examine if potential nonlinearities between fluorescence changes and the underlying spiking activity might affect our results, we used a nonlinear deconvolution method (MLspike; Deneux et al., 2016) to also estimate spiking. **(B)** MLSpike estimates with no onset nonlinearity (default parameters, MLspike; decay time constant of 0.4 seconds; amplitude parameter extracted via deconvolved estimates from OASIS; Methods). We computed the change in optogenetic responses as a function of visual responses. Pink line: LOESS fit with 95% confidence intervals via bootstrap. **(C)** MLSpike estimates with onset nonlinearity estimated from jGCaMP8s. MLspike estimation as above, with added onset nonlinearity estimation via polynomial fit; data from Zhang et al., 2023 (inset; black line: raw data; green line: fit polynomial; [β_0_, β_1_] = [-0.3, 0.06]). Purple line: LOESS fit with 95% confidence intervals. LOESS fits in **A**-**C** are cropped for visual comparison to the minimum range of visual responses spanned by the data across each of the 3 methods. **(D-E)** Deconvolved traces (mean ± SEM) using the constrained OASIS method for example neurons in Fig. 2F-M, showing attenuation in the visually-suppressed neuron.

**Fig. S6:**
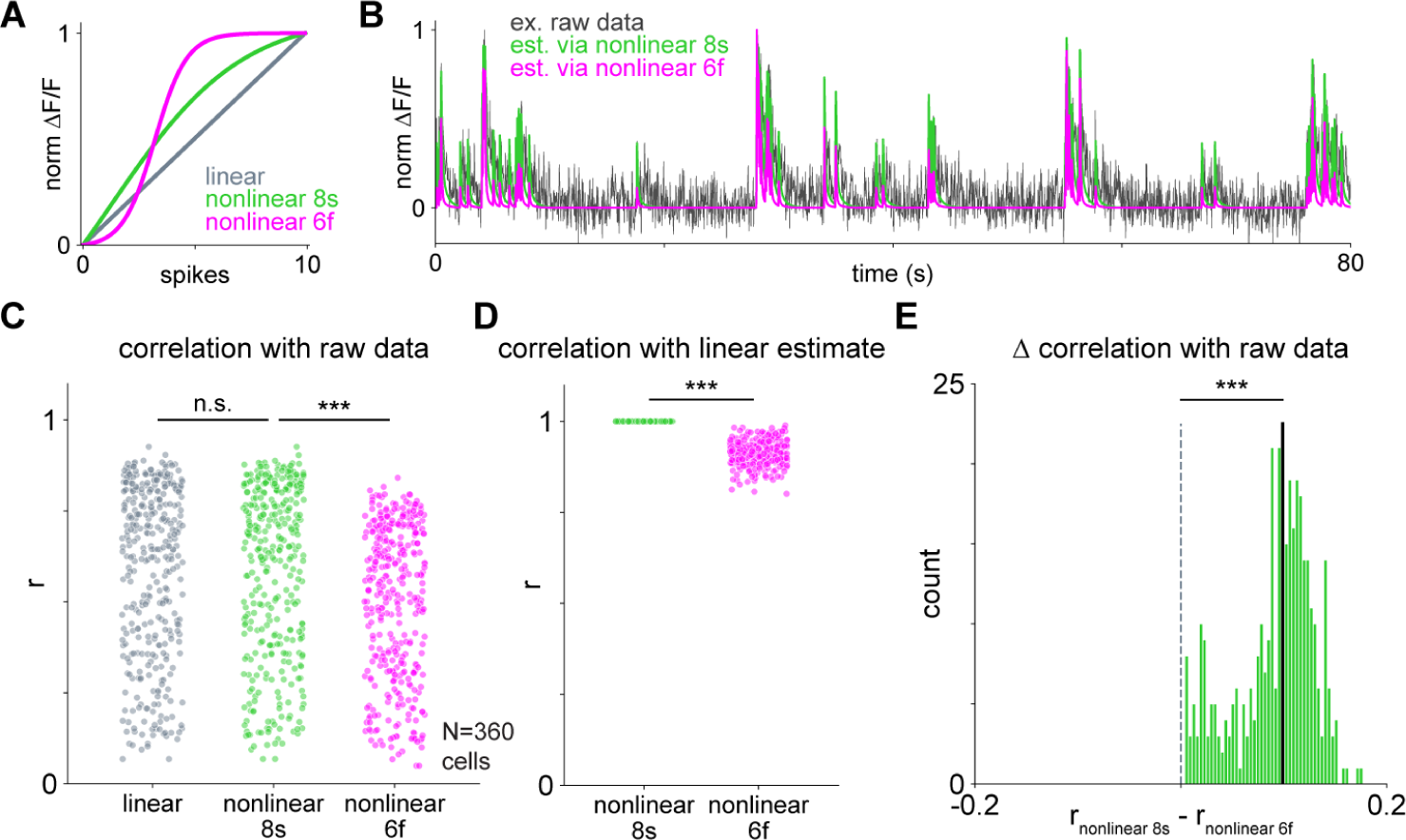
jGCaMP8s data is described by a nearly linear relationship between spikes and fluorescence. To examine if nonlinearities in the calcium indicator significantly impact our measurements, we passed spike time estimates generated by MLspike (nonlinear jGCaMP8s parameters; Fig. S5) into a double-decay spike-to-fluorescence (S2F) model that considers either nonlinear or linear relationships between spiking and fluorescence (Zhang et al., 2023). Using the nonlinearities measured in Zhang et al., 2023 for jGCaMP8s (the indicator used in our measurements) and for GCaMP6f (an indicator with a sharper nonlinearity for low spike count), as well as a linear function, we generated estimates of the calcium activity and compared them to the original calcium data. **(A)** The nonlinear (sigmoidal, for jGCaMP8s and GCaMP6f) and linear relationships between spiking and ΔF/F_0_. **(B)** Example calcium activity trace from raw data (gray) with estimated calcium traces using the nonlinear S2F model for 8s (green) and 6f (magenta). As expected, the GCaMP6f model does not describe the data well and the linear and jGCaMP8s model are similar. **(C)** Correlations (Pearson’s r) between the estimated calcium from the linear and nonlinear S2F models and the raw fluorescence data across cells (N = 360). The linear and nonlinear 8s models are similarly correlated to the raw data (p = 1.0; Mann-Whitney U test with Bonferroni correction), while the nonlinear 6f model shows smaller correlations with the raw data than the nonlinear 8s model (p < 0.001; Mann-Whitney U test with Bonferroni correction). **(D)** The nonlinear 8s model is well correlated to the linear model (mean correlation 0.99989, left), and the nonlinear 6f model performs less well (p < 0.001; Mann-Whitney U test) **(E)** Differences between the correlation of the raw data with either the nonlinear 8s or 6f models. The nonlinear 8s model provides a better estimate of the raw fluorescence for all neurons (p < 0.001; Wilcoxon signed-rank test). In sum, the relationship between spikes and fluorescence in our data is nearly linear, as also demonstrated in Zhang et al., 2023 for jGCaMP8s. The nonlinear 8s model is not a significant improvement over the linear model, supporting the use of OASIS for deconvolution analyses (Fig. S5).

**Fig. S7:**
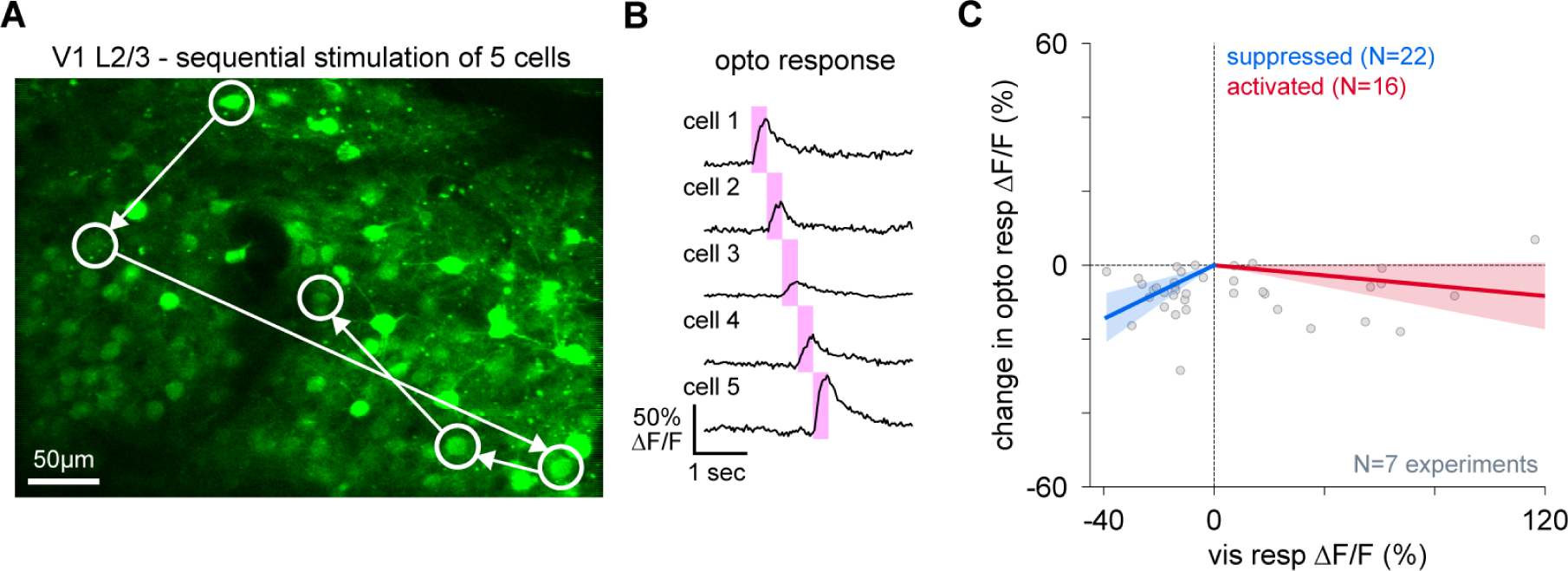
Optogenetic stimulation of individual neurons produces similar results to stimulation of small populations. **(A)** Example: targeting individual neurons in layer 2/3 of V1 expressing jGCaMP8s-P2A-stChrimsonR (N=5, each cell stimulated in sequence to improve experimental yield). **(B)** Example traces of sequential, non-overlapping stimulation in 5 example neurons. Pink regions: 300 ms optogenetic stimulation period. **(C)** Scatter plot of changes optogenetic responses as a function of visual response for individually-stimulated neurons (N = 7 experiments, N = 22 visually-suppressed cells, N = 16 visually-activated cells; visual stimulus: full-field gratings; cells below 5% ΔF/F response were excluded; Methods). Trial conditions same as in Figure 2 experiments. Lines: linear fits (intercept set to zero) with 95% confidence intervals for suppressed (slope = 0.36, R^2^ = 0.49, p = 2*10^-4^) and activated (slope = -0.07, R^2^ = 0.2, p = 0.07) cells. Here we used linear regression instead of LOESS fits due to smaller dataset size. However, these data show similar effects as the data with small populations stimulated: larger attenuation of responses during visual suppression and nearly linear effects during visual excitation.

**Fig. S8:**
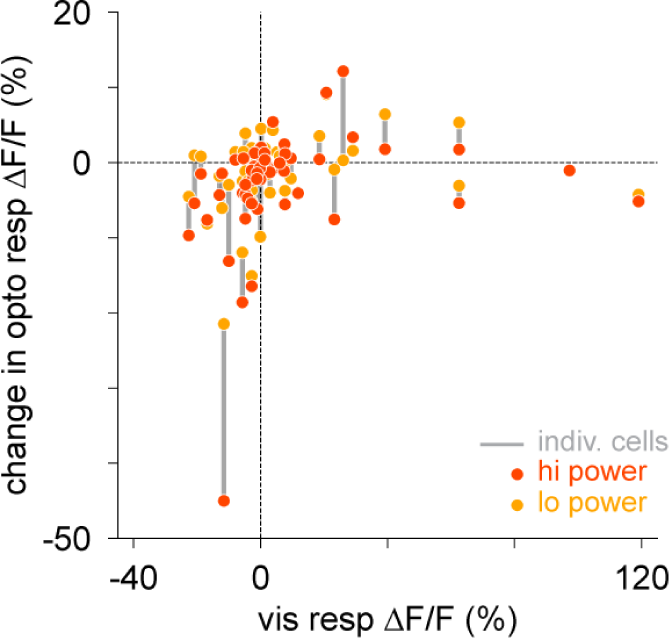
Changes in optogenetic responses during visual stimulation are similar for different optogenetic intensities. Change in optogenetic response as a function of average visual response measured using two optogenetic powers. Gray lines connect the two powers of stimulation for each individual cell.

**Fig. S9:**
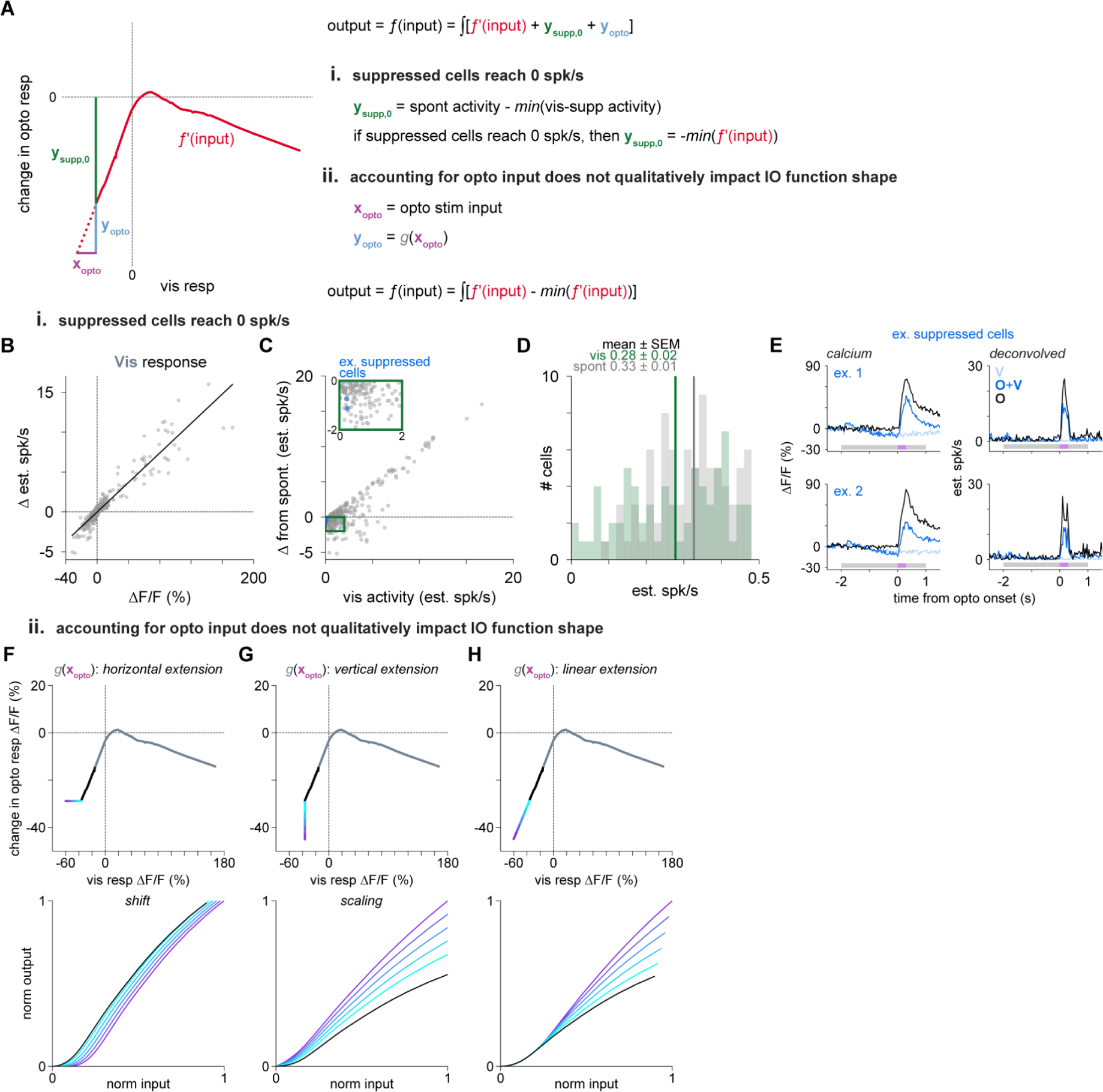
Estimates of the IO function do not depend on minimum firing rate. **(A)** Schematic showing how we measure the change in optogenetic response (change in output) as a function of visual response (input). This relationship is the derivative of the underlying IO function. Since the x-axis captures changes in response relative to baseline (spontaneous) activity, it is possible in principle that the choice of zero firing rate point could affect our results. Here we use deconvolution **(B-E)** to estimate the rate of calcium transients including the zero rate and show that this is consistent with our minimum fluorescence measurements reflecting low rates close to zero. We also **(F-H)** show the effects of different zero points and demonstrate that they do not qualitatively change our findings. Note that deconvolution analyses produce similarly shaped IO function estimates (Fig. S10) as the fluorescence-based measurements, further supporting the idea that we accurately capture low firing rates in our data. **(B)** Deconvolved spiking estimates of visual responses match corresponding ΔF/F_0_ values. **(C-D),** Deconvolved spiking estimates of visually-evoked activity. **(C)** Visually-suppressed cells reach 0 spk/s. Inset: zoom near zero visual activity. **(D)** Deconvolved cell activity reaches zero during visual stimulation but not during spontaneous activity, implying cells are suppressed to zero activity during IO measurements. **(E)** Trial-averaged activity from two example neurons near 0 spk/s during visual suppression (V). These cells are indicated by blue points in **C**. **(F-H)** Top: example effects of potential mismeasurement of zero points. The three panels (**F-H**, top) show three possible scenarios where potentially lower spike rates exist for visual responses (**F**), optogenetic responses (**G**), or both (**H**). We tested the impact of various extensions on our data to assess the qualitative impact on the estimated IO function. **(F)** Horizontal extensions shift the IO function. **(G)** Vertical extensions scale the IO function. **(H)** Linear extensions of the LOESS fit (estimated using least squares regression on the region where x < -15% ΔF/F_0_) do not impact nonlinearities at low input but scale saturating nonlinearities. In all three cases, our major effects — the supralinearity below spontaneous rates and the largely linear region for excited cells — remain present. Finally, direct estimation of the IO function on deconvolved data (Fig. S10) also produces similar IO function shapes. All these observations support the idea that our estimated IO function shapes are not modified by issues with measuring minimum firing rates.

**Fig. S10:**
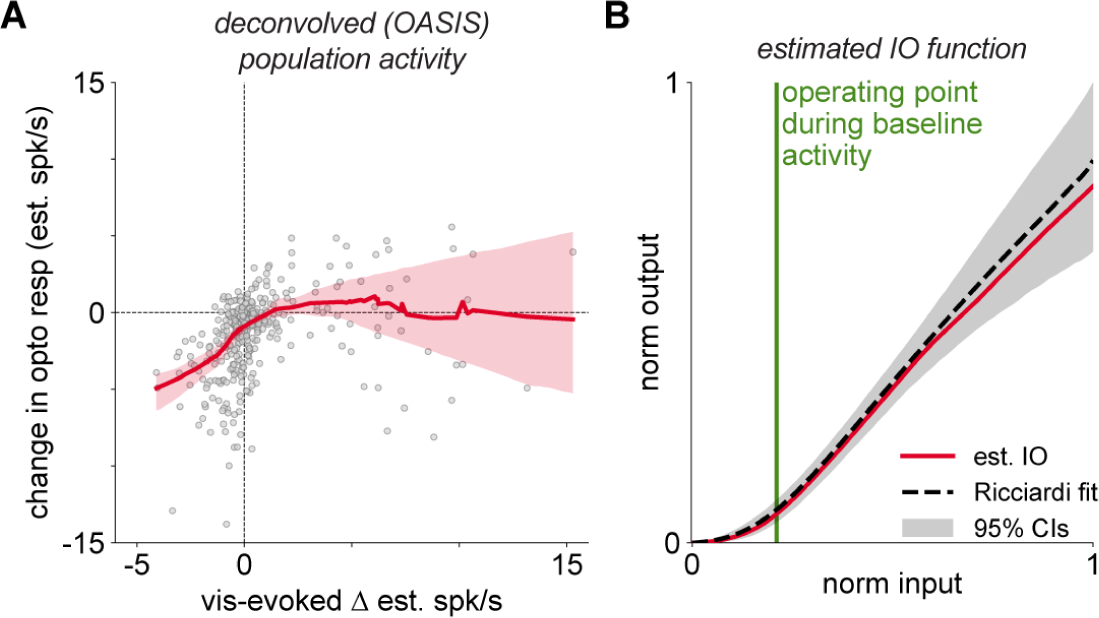
Deconvolved responses also show a supralinear-to-linear IO function, consistent with estimates from fluorescence. **(A)** Population (N = 364 cells) measure of change in deconvolved estimated spiking responses (OASIS) to optogenetic stimulation across the working range of visual responses. Red line: LOESS fit with 95% confidence intervals (via bootstrap), axis not rotated prior to fit. **(B)** Estimated IO function. Red: direct estimate from fit (integral of red line in A). Black dashed line, Ricciardi function fit, 95% confidence intervals (gray region; bootstrap), as in Fig. 4B. Green vertical line: operating point at rest (zero visual response in **A**).

**Fig. S11:**
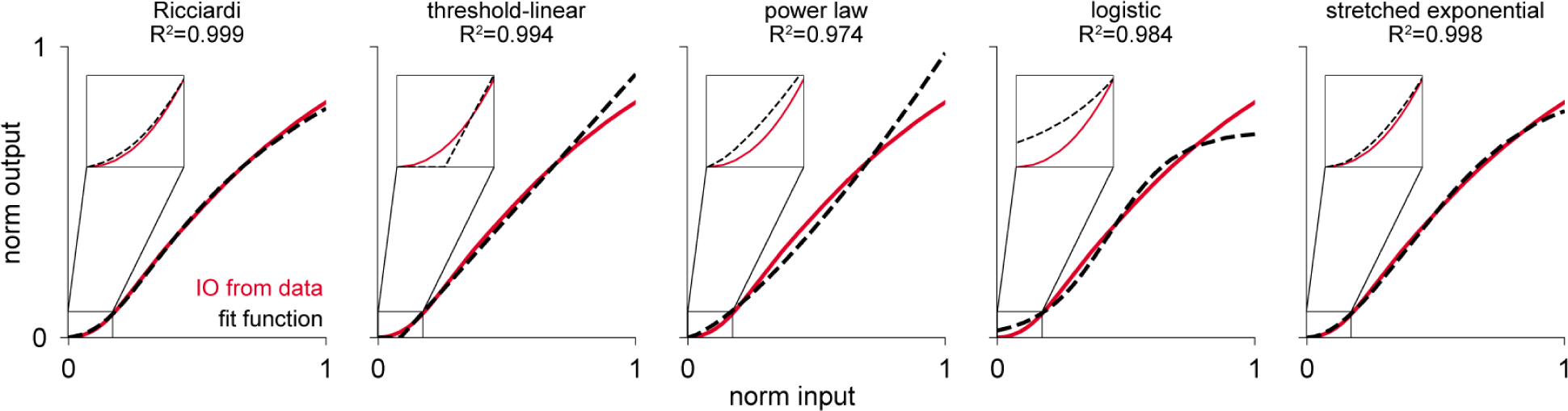
The estimated IO function is well-described by a Ricciardi function. Ricciardi, threshold-linear (rectified linear, or ReLU), power law, logistic, and stretched exponential function fits (black lines, least squares regression) to the estimated IO function (red line). Insets show comparison of curves at low inputs. Functions used for fits described in Methods. Ricciardi, logistic, and stretched exponential have three fit parameters, and threshold-linear and power law have two. The Ricciardi function is a good description of our data.

**Fig. S12:**
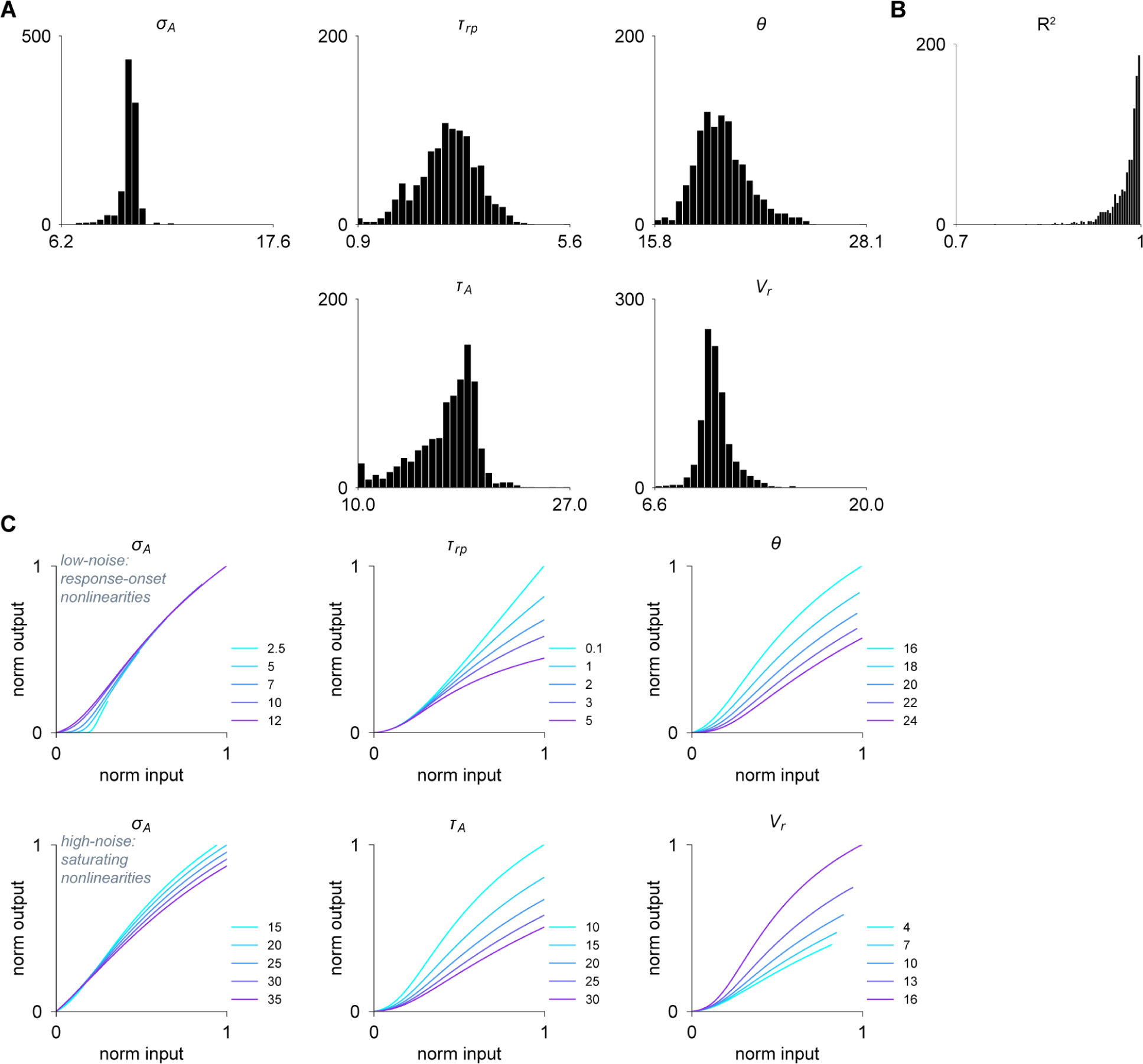
Ricciardi transfer function parameter sensitivity by bootstrap. **(A)** Distributions of parameters for bootstrapped Ricciardi function fit in Fig. 4B (N = 1000 repetitions; N = 364 samples). (**B**) Histogram of R^2^ fits of the Ricciardi functions (mean R^2^ = 0.975). **(C)** Ricciardi transfer function solutions for ranges of different parameters to show effects of each parameter. Changing parameter values across the range of bootstrap fits does not qualitatively change the shape of the transfer function solution. Parameters: *σ_A_* = 10, *τ_r_*_p_ = 2, *τ_A_* = 20, *θ* = 20, *V_r_* = 10, where not stated.

**Fig. S13:**
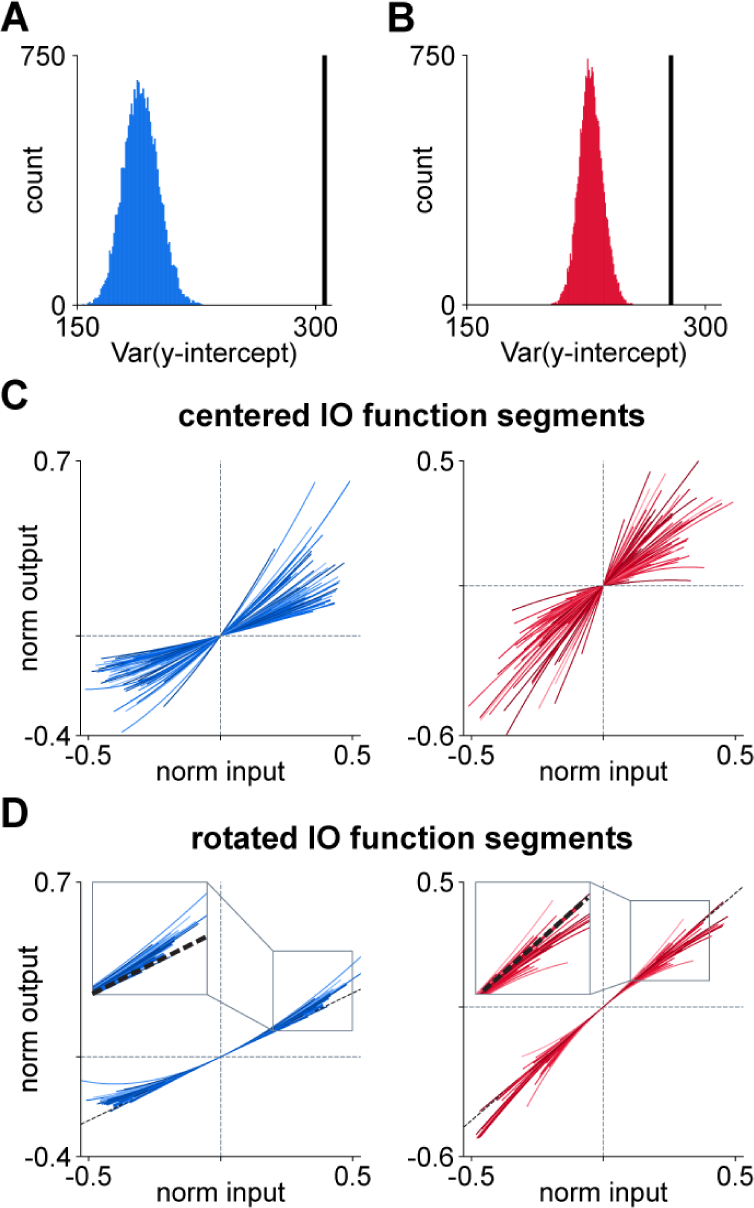
Cell-to-cell variance in IO function gain is not due to measurement noise. To assess if the variance across the fit intercepts (Fig. 5D) for cells from our mixed-effects model were greater than would be expected by chance due to data variability, we performed permutation tests by shuffling the preceding responses across all cells and all trials included for each cell (for either the suppressed or activated cells). **(A)** Histogram of distribution of variances in intercepts from the permutation test for suppressed cells. Black line: variance of the fit intercepts from Fig. 5D. p-value < 0.001. **(B)** Same as **A**, for activated cells. p-value < 0.001. **(C-D)** To visualize the variance between IO function slopes, we aligned the IO functions across cells (Fig. 5E) by first aligning their center point (**C**), then rotating each segment to match angle at the zero intercept. (**D**). Suppressed cells (blue, left) are supralinear: the majority fall above the black dashed line (slope at zero intercept), indicating a supralinear IO function. Activated cells (red, right) fall above and below the black line, showing that there is some variability in shape, but on net the activated IO function shape is linear.

